# Anionic lipids modulate mRNA-lipid nanoparticle immunogenicity and confer protection in a mouse model of multiple sclerosis

**DOI:** 10.1101/2025.10.17.683125

**Authors:** Jilian R. Melamed, Jenna Muscat-Rivera, Michael Kegel, Lesley S. Chaboub, Roxanne Perez-Tremble, Navdeep S. Bhalla, Houping Ni, Honghong Sun, Drew Weissman

**Affiliations:** Division of Infectious Diseases, Perelman School of Medicine, University of Pennsylvania, Philadelphia, PA 19104 USA; Penn Institute for RNA Innovation, Perelman School of Medicine, University of Pennsylvania, Philadelphia, PA 19104 USA; Department of Bioengineering, University of Pennsylvania, Philadelphia, PA 19104 USA; College of Arts and Science,, University of Pennsylvania, Philadelphia, PA 19104 USA; Department of Pathology and Lab Medicine, Perelman School of Medicine, University of Pennsylvania, Philadelphia, PA 19104 USA

**Author notes:** CORRESPONDING AUTHORS Drew Weissman, M.D., Ph.D., Roberts Family Professor in Vaccine Research, Director of the Penn Institute for RNA Innovation, Director of Vaccine Research in the Infectious Diseases Division Perelman School of Medicine, University of Pennsylvania 25 N 38th St, Philadelphia, PA 19104, USA, Jilian R. Melamed, Ph.D. Research Assistant Professor Perelman School of Medicine University of Pennsylvania, 25 N 38th St, Philadelphia, PA 19104, USA.

## Abstract

The modularity of mRNA-lipid nanoparticle (mRNA-LNP) platforms has enabled their rapid adaptation from infectious disease vaccines to emerging applications in immune-mediated disorders. However, extending mRNA-LNPs to autoimmune and inflammatory diseases requires precise control over immune cell targeting and immunogenicity. Here, we systematically investigate how incorporating anionic lipids into LNPs modulates both immune cell tropism and innate immune activation. Using a library of 40 distinct LNP formulations, we demonstrate that anionic lipids enhance mRNA delivery to splenic dendritic cells, reduce early cellular markers of adjuvant activity and tune cytokine responses in a lipid-dependent manner. We identify formulations that retain pro-inflammatory adjuvant activity and others that promote tolerogenic responses. A lead formulation containing the anionic lipid DOPG selectively dampens innate activation and induces IL-10 production. When encoding the myelin antigen MOG_35-55_, this LNP suppresses disease in a mouse model of multiple sclerosis, reducing neuroinflammation, T cell infiltration, and maintaining myelin morphology. These findings establish a framework for designing immune-targeted mRNA–LNPs with tunable immunogenicity and promote the development of antigen-specific tolerizing immunotherapies for autoimmune disease.

## Introduction

The rapid development and global deployment of mRNA-lipid nanoparticle (mRNA-LNP) vaccines during the SARS-CoV-2 pandemic has catalyzed widespread interest in extending this platform to a broader array of diseases driven by immune dysfunction. Central to this promise is the modularity of mRNA therapeutics: mRNA sequences can be engineered to encode virtually any protein, and LNPs can be chemically and biologically tuned to direct specific immune responses.^1–3^ This versatility positions mRNA-LNPs not only as vaccines for infectious diseases, but also as prophylactic and therapeutic agents for cancer, autoimmune disorders, allergies, gene therapy, and inflammatory conditions. Realizing this potential, however, requires a deeper mechanistic understanding of how LNPs can be rationally designed to target and modulate immune cell function.

Effective and safe delivery of mRNA–LNPs to immune cells is a prerequisite for immunological applications. While intramuscular administration exploits immune cells in draining lymph nodes to generate protective responses,^4^ many therapeutic contexts demand systemic delivery to diverse or disease-specific immune cell populations.^5,6^ Historically, systemic targeting of immune cells with mRNA–LNPs has been challenging, but recent advances have begun to overcome these barriers. For example, several ionizable lipids have demonstrated innate tropism for macrophages in lymphoid organs following intravenous administration.^7–10^ Yet, rational design of ionizable lipids remains difficult due to limited understanding of the structure–function relationships governing immune cell tropism.^11^

To address this, molecular targeting strategies have emerged, enabling cell-specific mRNA delivery independent of physicochemical tropism. Antibody-decorated LNPs targeting cell surface receptors have achieved efficient mRNA delivery to traditionally inaccessible populations such as T cells^12–14^ and hematopoietic stem cells.^6^ While powerful, this approach requires specialized manufacturing and relies on the availability of cell-specific surface markers that facilitate LNP internalization.

An alternative strategy, selective organ targeting (SORT), leverages the incorporation of anionic lipids to redirect mRNA–LNPs from the liver to lymphoid tissues.^15–18^ Unlike molecular targeting, SORT enhances mRNA delivery broadly to immune cells—including macrophages, dendritic cells, and lymphocytes—within the spleen and lymph nodes.^15–19^ The role of negative charge in immune cell targeting has also been observed in net-negative mRNA lipoplexes,^20–22^ which can elicit either cytotoxic CD8⁺ T cell responses^22^ or tolerogenic regulatory T cell (Treg) responses depending on the immunogenicity of the mRNA cargo.^21^ For instance, modified mRNA containing N1-methylpseudouridine can suppress innate immune activation and promote Treg induction,^21,23–25^ whereas unmodified mRNA can stimulate cytotoxic responses.^26^ This duality suggests that lipid charge and mRNA chemistry can be co-optimized to tune T cell responses and achieve tailored immunological outcomes.

Beyond mRNA modifications, LNP composition itself influences immune activation. Ionizable lipids, which become protonated in endosomes to facilitate mRNA release, also possess intrinsic adjuvant activity that promotes a Th1 biased T follicular helper cell and germinal center B cell responses.^27,28^ While beneficial for vaccines and cancer immunotherapy, this pro-inflammatory activity can be detrimental in contexts such as protein replacement or gene editing, particularly in patients with underlying inflammation.^11,29^ Thus, strategies to modulate LNP adjuvanticity are essential for expanding the therapeutic scope of mRNA–LNPs.

In this study, we sought to elucidate how LNP composition governs immune cell targeting and immunogenicity. We systematically evaluated a library of 40 LNP formulations varying in ionizable lipid structure, anionic phospholipid identity, and lipid molar ratios. Consistent with prior findings, we show that anionic lipids enhance mRNA delivery to splenic immune cells. We extend this knowledge by demonstrating that specific splenocyte populations, particularly macrophages and dendritic cells, can be preferentially targeted through lipid composition tuning.

Furthermore, we reveal that the nature and magnitude of immune responses elicited by mRNA–LNPs depend on both the ionizable and anionic lipid components. We identify formulations that retain adjuvant activity and others that promote tolerogenic responses. To validate the therapeutic potential of tolerogenic LNPs, we tested our lead formulation in a mouse model of multiple sclerosis (experimental autoimmune encephalomyelitis, EAE), where it reduced disease severity, increased Treg induction, and decreased spinal cord pathology.

Together, these findings provide mechanistic insights into the design of immune-targeted mRNA–LNPs and establish a framework for developing tailored immunotherapies for a wide range of diseases.

## Results and Discussion

Previous studies have demonstrated that anionic lipids can redirect LNP delivery to the spleen, a property with promising implications for immunotherapy. However, the influence of anionic phospholipids on the adjuvant activity of LNPs remains unexplored. Here, we systematically investigated how both ionizable and anionic phospholipids contribute to the immunostimulatory properties of LNPs, with the long-term goal of guiding the design of immune-tropic LNPs with tunable adjuvant activity. Such LNPs could be tailored for diverse therapeutic applications—for instance, highly immunogenic LNPs for cancer vaccines, and non-inflammatory LNPs for tolerogenic therapies targeting autoimmune diseases, allergies, or transplant rejection.

To this end, we generated a combinatorial library of 40 chemically distinct LNPs by varying three key parameters: (i) one of four ionizable lipids with known adjuvant profiles, (ii) one of three anionic phospholipids, and (iii) four molar ratios of lipid components (**Fig 1A**). The ionizable lipids SM-102 and ALC-0315 were selected for their demonstrated safety and efficacy in the SARS-CoV-2 vaccines and for their known Th1/Tfh adjuvant activity.^28,30,31^ MC3 was chosen because it is clinically approved for amyloidosis therapy^32^ and activates multiple inflammatory pathways,^33^ while we have previously demonstrated that 306O_i10_ is non-inflammatory and relatively versatile in its ability to deliver mRNA to multiple cell types.^34–36^ The anionic phospholipids DOPS, BMP, and DOPG were selected for their compatibility with standard LNP formulations.

**Figure 1.**
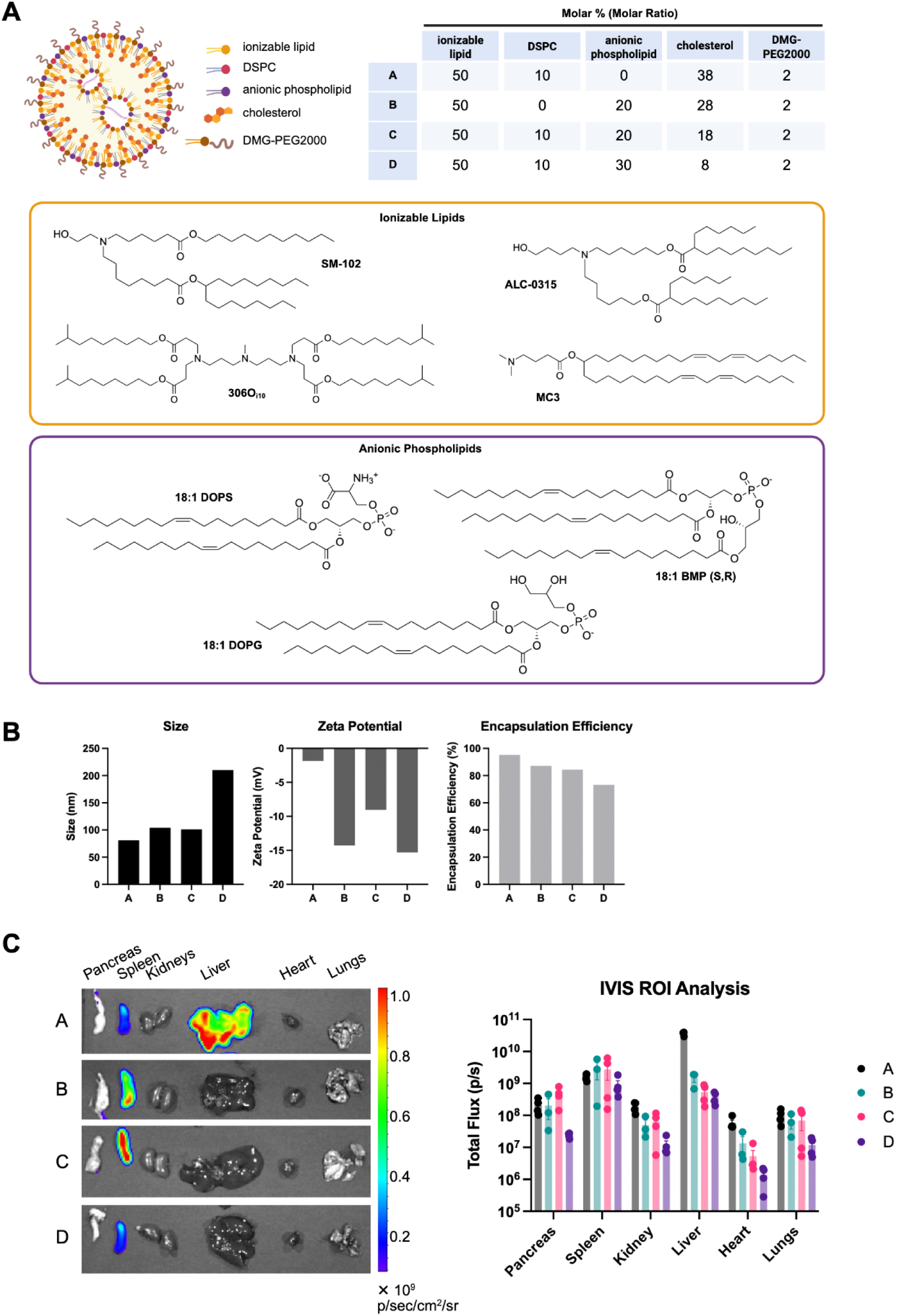
Combinatorial design and characterization of anionic lipid-containing mRNA–LNPs. ***A)*** Schematic of the 40-LNP library generated by varying ionizable lipid structure, anionic phospholipid identity (DOPS, BMP, DOPG), and molar ratios of lipid components. ***B)*** Physicochemical characterization of SM-102 LNPs with increasing DOPS content shows a DOPS-dependent increase in particle size, shift toward negative zeta potential, and reduced mRNA encapsulation efficiency. ***C)*** In vivo luciferase imaging 6 hours post-intravenous injection reveals that DOPS-containing LNPs redirect protein expression from the liver to the spleen, confirming splenic tropism of anionic SORT LNPs. Data shown are representative of 4 mice tested per group, with each point in the ROI quantification graph representing a single mouse and error bars show standard deviations.

We evaluated 4- and 5-component selective organ targeting (SORT)^15,16^ LNPs, with or without the zwitterionic “helper” DSPC, and with 0%, 20%, or 30% anionic lipid. Increasing DOPS content led to larger particle sizes: SM-102 LNPs without DOPS (composition A) measured ∼80 nm, while those with 20% DOPS (compositions B and C) increased to ∼100 nm, and those with 30% DOPS (composition D) reached ∼200 nm (**Fig 1B**). Zeta potential measurements revealed a shift toward greater net negative charge with increasing DOPS:DSPC ratios. Additionally, mRNA encapsulation efficiency declined with higher DOPS content, ranging from 95% (composition A) to 73% (composition D).

To validate that anionic lipids promote splenic delivery, we formulated SM-102 LNPs containing DOPS with mRNA encoding firefly luciferase (Luc), administered them intravenously (IV) to C57BL/6 mice (5 μg/mouse), and assessed protein expression 6 hours post-injection using an *in vivo* imaging system (IVIS). Consistent with prior reports on anionic SORT LNPs, DOPS decreased protein expression in the liver and enabled selective protein expression in the spleen (**Fig 1C**). The splenic tropism of anionic SORT LNPs has been well-documented by Dan Siegwart’s group, so we proceeded with our evaluation of adjuvant activity.^15,16^

### Anionic lipid-containing LNPs robustly transfect conventional type 2 dendritic cells in the spleen

To better understand the cellular mechanisms underlying the adjuvant activity of anionic SORT LNPs, we next sought to identify which splenic cell populations are transfected following systemic delivery. Since adjuvant activity is inherently linked to the engagement and activation of antigen-presenting cells (APCs), determining the biodistribution of mRNA expression across splenic immune subsets is critical. We hypothesized that anionic phospholipids not only redirect LNPs to the spleen but also influence cell-type specificity, thereby modulating the immunological outcome. To test this, we formulated our 40-LNP combinatorial library with mRNA encoding green fluorescent protein (GFP). 24 hours after IV administration to C57BL/6 mice, we harvested spleens and performed flow cytometry to quantify GFP expression across key immune populations in the spleen (**Fig 2A**), including dendritic cells (DCs), macrophages (Mφ), neutrophils (Neu), natural killer (NK) cells, B cells, and T cells. The gating scheme used to identify these cell populations is shown in **Fig 2B**. These data enabled us to correlate transfection efficiency with known roles in antigen presentation and cytokine production, providing mechanistic insight into how lipid composition governs LNP adjuvanticity.

**Figure 2.**
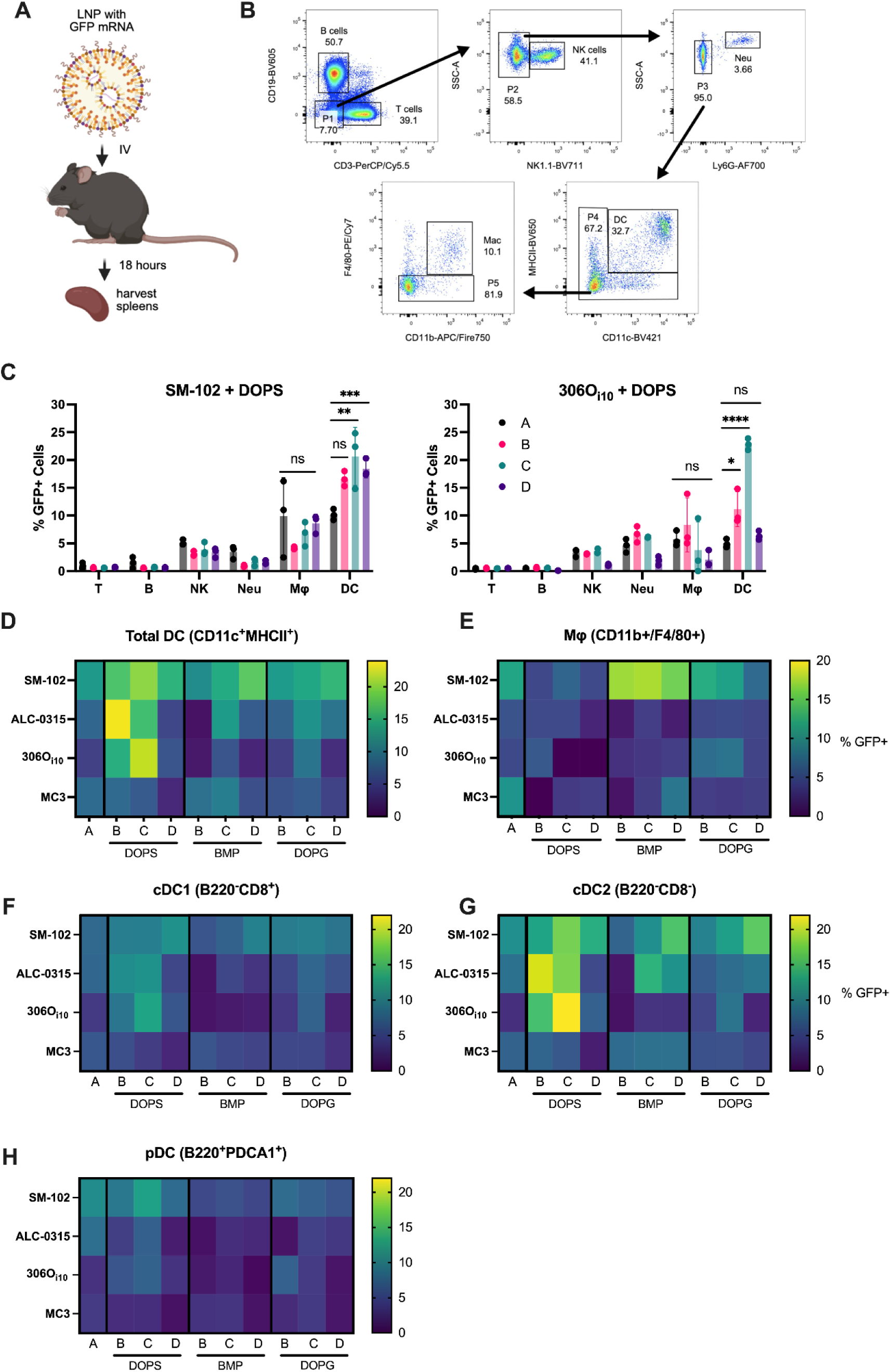
Anionic lipid-containing LNPs preferentially transfect antigen-presenting cells, particularly conventional type 2 dendritic cells (cDC2), in the spleen. *A)* Experimental workflow for assessing GFP expression in splenic immune cell populations following intravenous administration of GFP-encoding LNPs. *B)* Flow cytometry gating strategy used to identify major immune subsets. *C)* Representative transfection profiles for SM-102 and 306O_i10_ LNPs containing DOPS across splenic immune populations. Complete data for all groups are shown in **Fig S1**. *D-E)* Quantification of GFP⁺ dendritic cells and macrophages across formulations reveals that DOPS enhances DC transfection, while BMP selectively increases macrophage transfection. *F-H)* Subset analysis of dendritic cells shows preferential transfection of conventional type 2 DCs (cDC2), with minimal delivery to plasmacytoid DCs (pDCs) and conventional type 1 DCs (cDC1). The most effective formulations for cDC2 transfection were ALC-0315 with 20% DOPS and no DSPC, and 306O_i10_ with 20% DOPS and 10% DSPC. All experiments included n=3 mice/group, and data represent average measurements.

Across all formulations, LNPs most efficiently transfected macrophages and DCs, with minimal delivery to B and T cells and modest delivery to NK cells and neutrophils. The extent of transfection in APCs was strongly influenced by both the identity of the ionizable and anionic lipids, as well as their molar ratios. Representative breakdowns across all cell populations analyzed are shown for SM-102 and 306O_i10_ containing DOPS in **Fig 2C**; full data for all combinations are presented in **Fig S1**.

Among the anionic lipids tested, DOPS consistently enhanced delivery to DCs across all ionizable lipids except MC3, which showed poor transfection efficiency in splenocytes regardless of formulation. The most effective formulations for DC transfection were ALC-0315 with 20% DOPS and no DSPC and 306O_i10_ with 20% DOPS and 10% DSPC (**Fig 2D**). Notably, SM-102 LNPs containing BMP were the only formulations to markedly enhance GFP expression in macrophages (**Fig 2E**), suggesting that BMP may preferentially target this population or facilitate uptake via distinct mechanisms. BMP and DOPG also enhanced DC transfection when combined with SM-102, ALC-0315, and 306O_i10_, with the most pronounced effects observed in composition C, which included 20% anionic lipid and 10% DSPC.

Given the central role of DCs in initiating and shaping adaptive immune responses, we next sought to determine whether anionic SORT LNPs preferentially transfect specific DC subsets within the spleen. Conventional type 1 DCs (cDC1) specialize in cross-presentation and are critical for initiating cytotoxic T cell responses,^37,38^ while conventional type 2 DCs (cDC2) are adept at presenting antigens to CD4⁺ T cells and promoting T helper cell differentiation, particularly towards Th2 and T follicular helper (Tfh) phenotypes.^39^ In contrast, plasmacytoid DCs (pDCs), are key producers of type I interferons in response to viral stimuli.^40^ Characterizing the subset-specific tropism of LNPs is therefore critical to provide deeper mechanistic insight into how lipid composition influences adjuvant activity and immune polarization.

Across all formulations, we found that LNPs preferentially transfected cDC2 in the spleen, with minimal delivery to pDCs and cDC1 (**Fig 2F-H**). This pattern aligns with the Tfh-driven mechanism underlying protective immunity induced by mRNA-LNP vaccines.^27,28^ Consistent with our findings in total DCs (**Fig 2D**), the most effective formulations for cDC2 transfection were ALC-0315 with 20% DOPS and no DSPC and 306O_i10_ with 20% DOPS and 10% DSPC (**Fig 2G**). Notably, the latter formulation also achieved the highest transfection efficiency in cDC1. Although overall delivery to pDCs remained low, SM-102 with 20% DOPS and 10% DSPC maximized GFP expression in this population (**Fig 2H**). These results suggest that lipid composition can be tuned to selectively target DC subsets, offering a potential strategy to modulate immune polarization through rational LNP design.

### Anionic lipids attenuate LNP-induced dendritic cell maturation and T cell activation

The ability of LNPs to transfect dendritic cells is only one component of their immunological impact; the functional consequences of transfection, particularly DC maturation and subsequent T cell activation, are critical determinants of adjuvant activity. Having identified lipid compositions that selectively target DC subsets, we next investigated how anionic lipids influence the immunostimulatory potential of LNPs. Specifically, we assessed the extent to which these formulations promote DC maturation and drive downstream T cell responses, key hallmarks of effective stimulatory vaccine adjuvants.

DC maturation is characterized by the upregulation of costimulatory molecules that are essential for effective antigen presentation and T cell priming. In particular, CD40, CD80, and CD86 serve as canonical markers of DC activation, facilitating interactions with naïve T cells and promoting their differentiation (**Fig 3A**). CD80 and CD86 interact with CD28 on T cells to initiate lymphocyte proliferation and cytokine induction,^41,42^ while CD40 signals through its ligand, CD40L on activated T cells to increase the production of inflammatory cytokines such as IL-2 and IFNγ.^43,44^ Measuring the expression of these surface proteins therefore provides a direct readout of the immunostimulatory potential of each LNP formulation.

**Figure 3.**
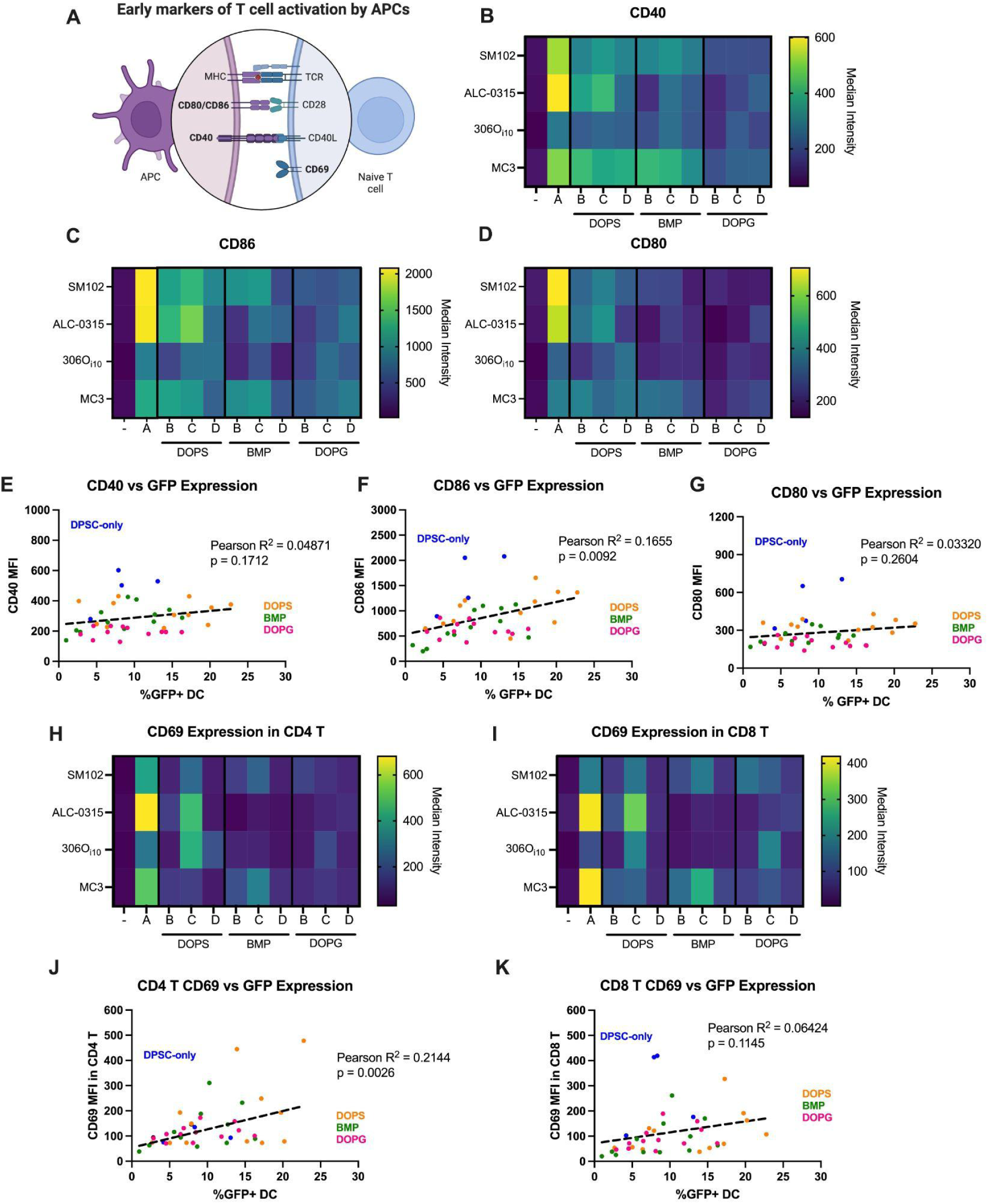
Anionic lipid-containing LNPs suppress dendritic cell maturation and early T cell activation. *A)* Schematic overview of dendritic cell (DC) maturation and T cell activation pathways, highlighting key costimulatory molecules. ***B-D)*** Flow cytometric analysis of CD40, CD86, and CD80 expression on splenic DCs 24 hours after intravenous administration of GFP-encoding LNPs reveals that incorporation of anionic lipids attenuates DC maturation. DOPG-containing formulations show the most pronounced suppression of CD40 and CD80 expression. ***E-G)***, Correlation between GFP⁺ DC frequency and costimulatory marker expression. DOPG-containing LNPs exhibit low activation despite efficient transfection. ***H-I)*** CD69 expression on CD4⁺ and CD8⁺ T cells 24 hours post-injection shows reduced early T cell activation in mice treated with anionic lipid-containing LNPs, particularly those formulated with BMP and DOPG. ***J-K)***, Correlation between GFP⁺ DCs and CD69⁺ T cells reveals a significant association for CD4⁺ but not CD8⁺ T cells, suggesting that lipid composition can modulate DC–T cell crosstalk and downstream immune activation. All experiments included n=3 mice/group, and data represent average measurements. Complete data are shown in **Fig S2**.

Using flow cytometry to measure the expression of cellular activation markers 24 hours after IV administration of mRNA-LNPs, we found that incorporating anionic phospholipids into otherwise inflammatory LNPs markedly suppressed DC maturation. The upregulation of CD40 (**Fig 3B**), CD86 (**Fig 3C**), and CD80 (**Fig 3D**) was significantly reduced across most formulations containing anionic lipids (**Fig S2**). LNPs formulated with SM-102 or ALC-0315 in the absence of anionic phospholipids induced robust expression of these costimulatory molecules, consistent with their efficacy as vaccine adjuvants. However, the addition of any anionic lipid, regardless of identity or molar ratio, substantially diminished DC activation, with DOPG generally producing the lowest levels of CD40 and CD80 expression among LNP-treated groups. 306O_i10_ was the least stimulatory overall, though formulations containing BMP or DOPG still led to further reductions in CD86 and CD80 expression. MC3 induced relatively low levels of DC maturation compared to SM-102 and ALC-0315, and these levels remained consistent across anionic lipid types and compositions, likely reflecting its poor transfection efficiency in splenic DCs.

To disentangle the effects of lipid composition on DC activation from its effects on mRNA delivery, we next sought to correlate the expression of CD40, CD80, and CD86 with GFP fluorescence (**Fig 3E-G**). This approach allowed us to determine whether LNP-induced maturation is a direct consequence of transfection, or whether certain lipid formulations activate DCs independently of mRNA delivery. While overall correlations between activation markers and the percentage of GFP⁺ DCs were modest, CD86 expression showed a statistically significant association with GFP levels (**Fig 3F**), suggesting a partial link between transfection and activation. From this analysis, we would suggest that LNPs characteristically above the correlation line, mostly DSPC-only or DOPS-containing LNPs, are likely more inclined to retain their adjuvant activity, whereas those below the line, primarily DOPG-containing LNPs, exhibited reduced maturation despite comparable transfection. These findings suggest that certain lipid compositions may decouple mRNA delivery from immunostimulation, offering a strategy to fine-tune LNPs for either immunogenic or tolerogenic applications.

Our observed downregulation of the costimulatory molecules CD40, CD86, and CD80, while maintaining efficient mRNA delivery to dendritic cells (DCs), is particularly noteworthy given the central role of these markers in T cell activation. Their suppression is a critical prerequisite for inducing antigen-specific immune tolerance. This feature holds promise for the development of novel therapies targeting autoimmune diseases and allergies, which are driven by dysregulated immune recognition. Supporting this concept, several preclinical studies have demonstrated that silencing CD40, CD86, and CD80 using antisense oligonucleotides or siRNA can promote antigen-specific tolerance in models of autoimmunity, allergy, and transplant acceptance.^45–48^ Similarly, Krienke et al. reported that a liposomal formulation of nucleoside-modified, autoantigen-encoding mRNA attenuated the expression of costimulatory molecules compared to unmodified mRNA, thereby restoring immune tolerance in a murine model of multiple sclerosis.^21^

To further evaluate the downstream immunological consequences of LNP-induced dendritic cell activation, we next examined early T cell responses. Activation of naïve T cells requires not only antigen presentation but also costimulatory signals from mature DCs. One of the earliest markers of T cell activation is CD69, a surface protein rapidly upregulated following T cell receptor engagement. We therefore assessed CD69 expression on CD4⁺ and CD8⁺ T cells 24 hours after LNP administration, aiming to correlate T cell activation with the maturation status and transfection efficiency of upstream DCs.

Consistent with our observations of DC maturation markers, we found that LNPs lacking anionic lipids elicited the strongest upregulation of CD69 expression in both CD4⁺ and CD8⁺ T cells, with ALC-0315 and MC3 demonstrating particularly robust induction (Fig 3 **H,I**). In contrast, SM-102 and 306Oi10 without anionic lipids induced only moderate CD69 expression. Incorporating anionic helper lipids generally attenuated CD69 upregulation across both T cell subsets. However, formulations containing 10% DSPC and 20% DOPS maintained relatively high levels of CD69 induction. Notably, removal of DSPC or increasing the DOPS content to 30% suppressed CD69 upregulation, suggesting that the balance of helper phospholipid charge plays a critical role in modulating LNP adjuvanticity. The effect appeared unique to DOPS, as the other anionic phospholipids, BMP and DOPG, enabled little CD69 expression at any composition ratio.

To better understand how lipid composition influences T cell activation independently of its impact on mRNA delivery efficiency, we next analyzed the relationship between GFP expression in DCs and CD69 expression in T cells (**Fig 3J,K**). We observed a significant positive correlation between DC transfection efficiency and CD69 expression in CD4⁺ T cells (**Fig 3J**), consistent with previous studies showing that LNPs preferentially elicit strong CD4⁺ T cell responses.^27,28^ In contrast, no significant correlation was detected between GFP expression in DCs and CD69 levels in CD8⁺ T cells (**Fig 3K**). Notably, formulations that deviated below the correlation trendline, indicative of reduced adjuvant activity, primarily contained the anionic lipids DOPS or DOPG.

These results demonstrate that SORT LNPs containing anionic lipids can be tuned to selectively suppress DC maturation and downstream T cell activation independently of achieving robust mRNA delivery to DCs. This decoupling of transfection from immunostimulation provides a modular platform for designing LNPs with tailored immunological profiles, enabling applications that span both vaccine development and immune tolerance induction.

### Anionic lipids suppress pro-inflammatory cytokine induction by LNPs

Building on our findings that anionic lipids attenuate DC maturation and early T cell activation markers, we next investigated the systemic cytokine responses elicited by mRNA-LNP administration. Cytokines serve as key mediators of innate immunity and are often used as biomarkers of vaccine reactogenicity and adjuvant potency. Because LNP-induced cytokine release can influence both the magnitude and quality of adaptive immune responses, we sought to determine how lipid composition modulates the inflammatory milieu following mRNA delivery. By profiling serum cytokines 6 hours post-injection, we aimed to capture the acute innate immune signature associated with each formulation and assess its alignment with observed cellular activation patterns.

We first assessed cytokine induction across the four ionizable lipids formulated into conventional LNPs without anionic helper lipids (**Fig 4A**). SM-102 and ALC-0315, consistent with their known Th1/Tfh-skewing adjuvant activity, induced robust levels of IL-6 and IL-18. MC3 triggered elevated levels of IL-6, TNFα, and IL-5 (**Fig 4A,F**), whereas 306O_i10_ did not elicit detectable levels of any measured cytokines.

**Figure 4.**
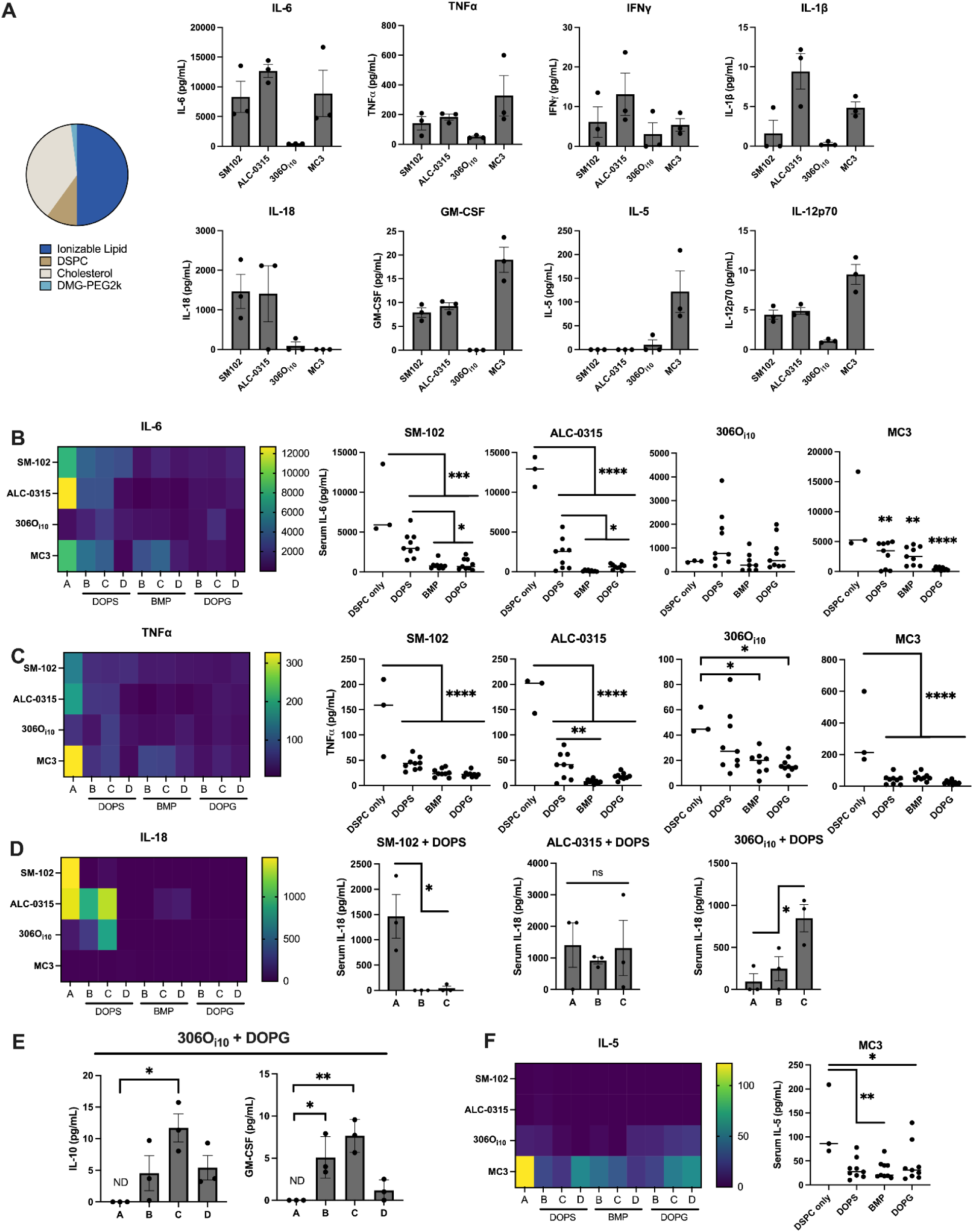
Anionic lipid-containing LNPs suppress pro-inflammatory cytokine responses following mRNA delivery. *A)* Serum cytokine levels 6 h after intravenous administration of mRNA–LNPs formulated with four ionizable lipids and no anionic phospholipids. SM-102 and ALC-0315 induced robust IL-6 and IL-18; MC3 elicited IL-6, TNFα, and IL-5; 306Oi10 showed minimal cytokine induction. ***B-D)*** Inclusion of anionic phospholipids (DOPS, BMP, DOPG) reduced cytokine levels across formulations, with DOPG showing the strongest suppression. ***E)*** DOPG-containing 306O_i10_ LNPs uniquely induced IL-10 without accompanying pro-inflammatory cytokines, suggesting potential for anti-inflammatory signaling. ***F)*** Only MC3-based LNP induced Th2 cytokines such as IL-5, which was already reduced with anionic phospholipids. These data demonstrate that anionic lipids can attenuate LNP adjuvanticity and modulate systemic cytokine signatures. In all graphs, individual data points represent individual mice. Error bars represent standard deviations. Statistical differences were calculated by ANOVA with post-hoc Tukey test. *p<0.05, **p<0.01, ***p<0.005, ****p<0.001.

We next evaluated the impact of anionic phospholipids on cytokine induction by LNPs (**Fig 4B-D, Fig S3**). For all ionizable lipids, we generally observed decreases in all cytokines in anionic lipid-containing formulations compared with their anionic lipid-free analogs. Among these, DOPS-containing LNPs retained the most adjuvant activity, with SM-102 and ALC-0315 DOPS formulations inducing significantly more IL-6 than those containing BMP or DOPG (**Fig 4B**). Interestingly, DOPS-containing 306Oi10 LNPs induced high levels of IL-18 (**Fig 4D**), unlike the DSPC-only LNP, though this was not accompanied by IL-6 (**Fig 4B**). Of all formulations tested, 306O_i10_ LNPs containing DOPG were the only ones to elicit detectable levels of IL-10 without the accompaniment of other inflammatory cytokines, suggesting a potential for anti-inflammatory signaling (**Fig 4E, Fig S3**).

MC3-based LNPs elicited the broadest range of cytokine production, including Th1, Th2, and Treg-skewing cytokines (**Fig 4A-C,F, Fig S3**). Notably, anionic lipid incorporation generally decreased all of these cytokines, with the greatest effects observed for IL-5 and IL-6. Similar trends were observed for GM-CSF, IL-12p70, IL-4, and IL-10, though their levels overall remained low (**Fig S3**).

Together, these findings demonstrate that anionic lipid incorporation can selectively dampen pro-inflammatory cytokine responses to mRNA–LNPs, offering a tunable strategy to modulate innate immune activation.

### Anionic lipid-containing LNPs reduce disease severity in a mouse model of multiple sclerosis

Having established that anionic lipids modulate the innate immune profile of mRNA–LNPs, we next sought to evaluate the functional consequences of these immunomodulatory effects *in vivo*. Given their reduced adjuvant activity, we hypothesized that anionic lipid-containing LNPs might mitigate aberrant immune responses in disease settings. To test this, we employed the experimental autoimmune encephalomyelitis (EAE) model, a well-established murine model of multiple sclerosis (MS),^49^ to investigate how LNP lipid composition influences therapeutic efficacy and immune polarization in the context of autoimmune neuroinflammation.

MS is a chronic autoimmune disorder of the central nervous system (CNS) characterized by immune-mediated destruction of myelin, the protective sheath surrounding neurons.^50^ This demyelination is driven by infiltration of autoreactive T and B cells and activation of resident glial cells, including astrocytes and microglia.^51,52^ The resulting neuroinflammation leads to progressive deficits in motor coordination, vision, and balance. MS remains one of the leading causes of neurological disability in young adults.^50^ Despite advances in disease-modifying therapies, current treatments rely on broad immunosuppression, which often leads to systemic side effects and increased infection risk, underscoring the need for more targeted approaches.^53^

In contrast, antigen-specific tolerizing immunotherapies aim to selectively silence pathogenic immune responses while preserving global immune competence. Delivering disease-relevant antigens in a non-inflammatory context has the potential to re-establish immune tolerance and halt disease progression without the systemic side effects associated with broad immunosuppression. In this context, the ability of anionic lipid-containing LNPs to dampen innate immune activation while enabling targeted antigen delivery positions them as a promising platform for next-generation MS therapies.

To evaluate this potential, we selected DOPG as the anionic lipid based on its superior ability to blunt adjuvant activity and its potential to induce IL-10 production. We incorporated DOPG into LNPs containing the ionizable lipid 4A3-SC8, which is structurally similar to 306O_i10_ but with the addition of thioether groups in its alkyl chains, facilitating enhanced biodegradability and safety. The composition of these LNPs is shown in **Fig 5A**.

**Figure 5.**
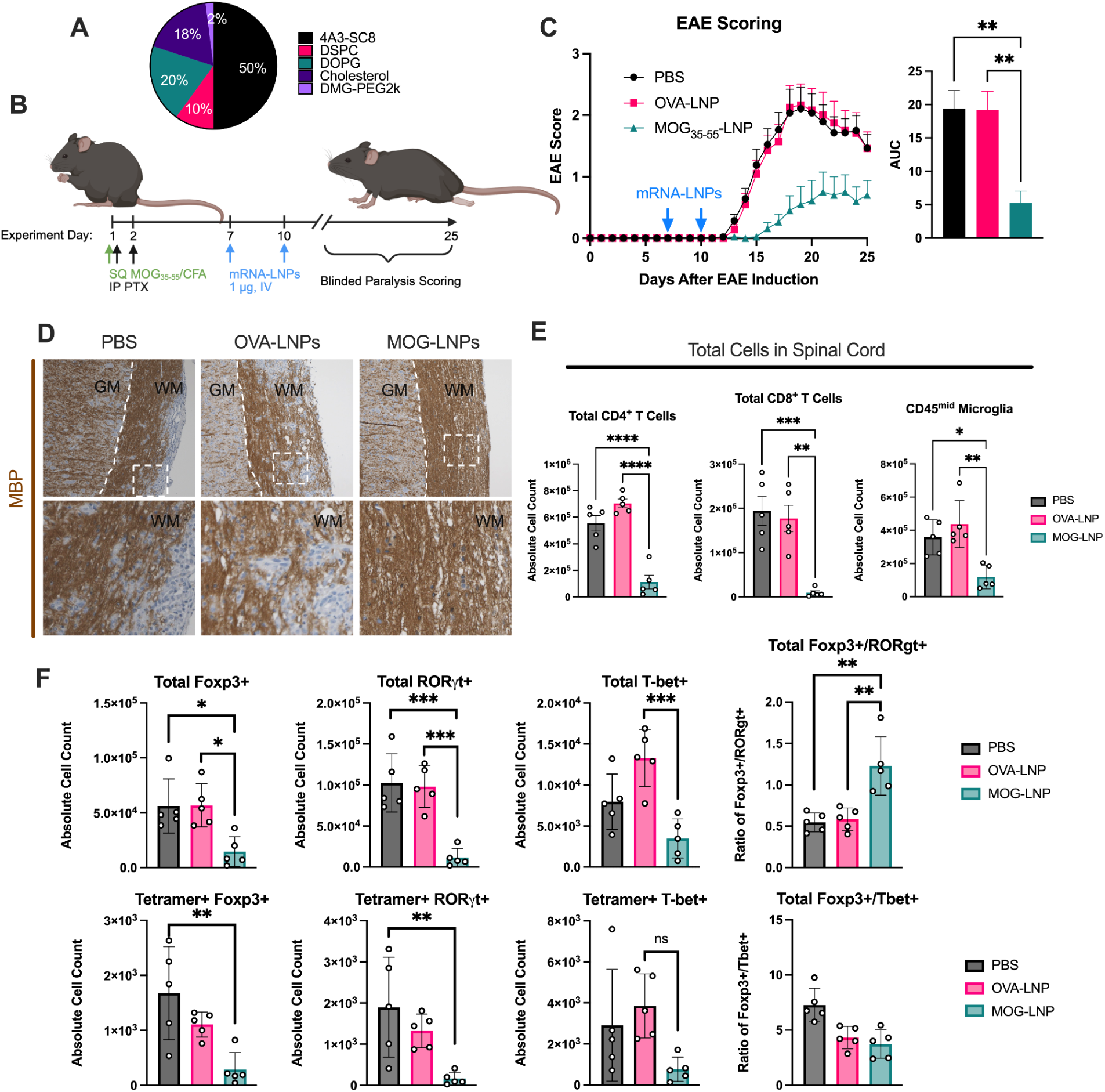
Anionic lipid-containing LNPs suppress neuroinflammation and reduce disease severity in a mouse model of multiple sclerosis. *A)* Composition of the lead tolerogenic LNP formulation containing the ionizable lipid 4A3-SC8 and the anionic lipid DOPG. *B)* Experimental timeline for EAE induction and mRNA-LNP treatment. *C)* Clinical scores over time show that MOG_35-55_-LNPs significantly delay disease onset and reduce paralysis severity compared to OVA-LNP and PBS controls. Data shown represent n>10 mice/group. *D)* Histological analysis of spinal cords at peak disease (day 18) reveals preserved myelin integrity and reduced immune cell infiltration in MOG_35-55_-LNP-treated mice. Dashed white lines visually separate gray and white matter. White dashed boxes indicate the enlarged region shown below. Scale bar = 100 μm. *E)* Flow cytometry of cells isolated from spinal cords shows reduced infiltration of CD4⁺ and CD8⁺ T cells and microglia in MOG_35-55_-LNP-treated mice. *F)* Tetramer and transcription factor staining of spinal cord CD4⁺ T cells demonstrates reduced MOG_35-55_-specific T cell infiltration and a shift toward a regulatory phenotype, with increased Foxp3⁺:RORγt⁺ ratios. These data support the therapeutic potential of tolerogenic mRNA-LNPs for antigen-specific immune modulation in autoimmune neuroinflammation.

EAE was induced in 10-week-old female C57BL/6 mice via subcutaneous (SQ) immunization with the myelin-derived peptide myelin oligodendrocyte glycoprotein 35-55 (MOG_35-55_) emulsified in Complete Freund’s Adjuvant (CFA).^54^ Pertussis toxin (PTX) was administered intraperitoneally at 3 and 24 hours post-immunization to transiently disrupt the blood–brain barrier and facilitate CNS infiltration by autoreactive lymphocytes. While human MS is mediated by complex immune networks, EAE is primarily mediated by Th1- and Th17-polarized CD4+ T cells.^55^

To assess the therapeutic efficacy of DOPG-containing LNPs, mice received intravenous injections of LNPs encoding either MOG_35-55_ or an irrelevant control antigen (OVA), or PBS on days 7 and 10 post-induction (1 μg mRNA per dose). Disease progression was monitored by blinded observers using a standardized EAE clinical scoring system through day 25 (**Fig 5B, Fig S4**). Mice treated with MOG_35-55_-encoding LNPs exhibited a delay in symptom onset and a significant decrease in disease severity compared with both OVA-LNP and PBS-treated controls (**Fig 5C**). These results demonstrate that immunomodulatory, anionic lipid-containing LNPs can deliver tolerogenic antigens *in vivo* to suppress autoimmune pathology, supporting their potential as a platform for antigen-specific immunotherapy in MS and other T cell-mediated autoimmune diseases.

To better understand the mechanism underlying the observed clinical benefit, we next examined immune cell populations within the CNS. Given that EAE is driven by infiltration of autoreactive lymphocytes in the spinal cord, we performed histological and flow cytometric analyses to assess how mRNA-LNP treatment influenced immune cell composition and activation at the site of inflammation. At the peak of disease (day 18), mice were sacrificed, perfused, and spinal cords were harvested for histological evaluation. Tissue sections were stained for myelin basic protein (MBP) to assess demyelination and immune infiltration. PBS- and OVA-LNP–treated mice exhibited extensive regions of demyelination accompanied by hypercellularity, indicative of robust immune infiltration (**Fig 5D**). In contrast, spinal cords from MOG_35-55_-LNP–treated mice retained morphologically normal myelin, as evidenced by uninterrupted MBP staining in the white matter with minimal cellular infiltration (**Fig 5D**). These histological findings closely mirrored clinical scores, with reduced demyelination corresponding to lower paralysis severity in MOG_35-55_-LNP–treated animals.

Flow cytometric analyses supported the histological observations. Spinal cords were dissected from EAE-induced mice at peak disease (day 18), enzymatically digested into single-cell suspensions, and analyzed for immune cell populations. Mice treated with MOG_35-55_-LNPs exhibited significantly fewer infiltrating CD4⁺ and CD8⁺ T cells compared to PBS and OVA-LNP controls (**Fig 5E**), consistent with reduced neuroinflammation. We also observed a reduction in microglia, the resident immune cells of the CNS, in MOG_35-55_-LNP–treated mice (**Fig 5E**). In MS, microglia play a dual role: they are essential for clearing myelin debris and supporting tissue repair, but their overactivation contributes to a pro-inflammatory environment that impairs remyelination. The reduced microglial presence observed in MOG_35-55_-LNP–treated mice may reflect a lower level of local immune activation, which could be critical for enabling regenerative processes within the CNS. Together, these findings suggest that anionic lipid-containing LNPs not only limit peripheral T cell infiltration but also dampen local innate immune activation within the CNS.

To further characterize the phenotype and antigen specificity of infiltrating CD4⁺ T cells, we performed transcription factor and tetramer staining on spinal cord–derived cells. We assessed expression of Foxp3, RORγt, and T-bet, key transcription factors that define Tregs, Th17 cells, and Th1 cells, respectively. Mice treated with MOG_35-55_-LNP exhibited a marked reduction in both total and MOG_35-55_ tetramer-positive CD4⁺ T cells within the spinal cord, indicating a suppression of antigen-specific effector T cell infiltration (**Fig 5F**). Notably, the ratio of Foxp3⁺ to RORγt⁺ cells was significantly increased in MOG_35-55_-LNP–treated mice, suggesting a shift in the local immune environment toward a more regulatory, less inflammatory phenotype. This skewing toward tolerance is consistent with the observed clinical protection and supports the potential of tolerogenic LNPs to reprogram autoreactive T cell responses in situ (**Fig 5F**).

To assess the potential for systemic toxicity associated with LNP treatment during EAE, we collected serum and organs from mice on day 18 post-induction, eight days after the final mRNA-LNP administration, when inflammatory burden was expected to peak. All animals exhibited signs of systemic inflammation, including detectable serum levels of IFNγ, IL-6, IL-10, and IL-27 (**Fig S5A**), and hypercellularity in the lungs and spleen (**Fig S5B**), unsurprising for mice at the peak of EAE severity. However, none of these inflammatory markers were significantly elevated in LNP-treated mice compared to PBS controls. These findings suggest that tolerogenic LNPs do not exacerbate systemic inflammation during autoimmune disease progression, supporting their safety profile for therapeutic use in chronic inflammatory settings.

Together, these results demonstrate that LNPs containing certain anionic and ionizable lipids encoding autoantigens can effectively suppress neuroinflammation, reduce pathogenic T cell infiltration, and promote a regulatory immune environment within the CNS. By incorporating anionic lipids to tune the adjuvant properties of LNPs, this platform offers a promising strategy for re-establishing immune homeostasis in autoimmune disease and other inflammatory disorders in an antigen-specific manner.

Extensive research has shown that non-inflammatory delivery of autoantigens can elicit antigen-specific immune tolerance in models of MS,^56–61^ type 1 diabetes,^62–66^ Celiac disease,^67^ and rheumatoid arthritis.^68,69^ While such protein-based approaches have undoubtedly shown efficacy and have progressed to clinical trials,^5^ they are often limited by manufacturing complexity, poor solubility, and delivery challenges. Nanoparticle encapsulation can improve delivery, but each autoantigen typically requires a custom formulation, reducing scalability and modularity. This is particularly problematic in human autoimmune diseases, which often involve multiple autoantigens. Additionally, many tolerogenic peptides are constrained by HLA restriction, limiting their applicability across diverse patient populations. In contrast, mRNA-based approaches overcome these limitations by enabling endogenous antigen expression, modular design, and compatibility with standardized delivery platforms. The rapid production process for new mRNA sequences also poises this platform to enable personalized tolerizing therapies tailored to individual autoimmune profiles and biomarkers.

However, the intrinsic adjuvant activity of LNPs has historically limited their application in tolerizing therapies.^27,28^ This pro-inflammatory signaling can exacerbate pre-existing inflammation,^29^ posing additional challenges for treating autoimmune diseases, which are characterized by chronic immune activation.^11,70^ Nonetheless, recent advances in mRNA-LNP design have enabled the development of non-inflammatory formulations capable of delivering antigens in a tolerogenic context. For instance, BioNTech developed a liposomal platform with a net negative charge balance that enhances targeting to splenic immune cells.^20^ When formulated with nucleoside-modified mRNA, these lipoplexes enabled non-inflammatory autoantigen delivery, resulting in delayed onset and reduced EAE severity in mice.^21^ Critically, this platform lacked an ionizable lipid, which contributed to its tolerogenic profile but required relatively high mRNA doses (two doses of 40 μg per mouse) to achieve therapeutic efficacy that is not translatable to human therapy. In contrast, LNPs leveraging non-inflammatory ionizable lipids and incorporating anionic helper lipids are able to achieve protection from EAE using substantially lower doses, as we have demonstrated here with two doses of 1 μg per mouse and others have demonstrated with DOPS-containing LNPs.^71^

Negatively charged nanoparticles are frequently employed to dampen inflammation and promote tolerogenic immune responses.^11,70,72^ One proposed mechanism is that their uptake and potential recycling by antigen-presenting cells mimics the clearance of apoptotic cells, an inherently tolerogenic process.^73,74^ This has been well-documented for phosphatidylserine (PS)-containing liposomes and LNPs,^71,75,76^ as PS is naturally exposed on the outer leaflet of apoptotic cell membranes. Interestingly, similar tolerogenic effects have been observed with liposomes and LNPs containing phosphoglycerol (PG) lipids,^77^ suggesting that the immunological consequences of negative surface charge may extend beyond PS-specific signaling. Interestingly, our data suggest that DOPG may elicit even stronger tolerogenic responses than DOPS-containing counterparts, highlighting PG as a promising alternative for designing non-inflammatory mRNA delivery systems.

In summary, our findings demonstrate that incorporating anionic phospholipids into LNPs provides a strategy not only to target desired immune cell populations but also to tailor the adjuvant activity of LNPs for a given application. By leveraging this design flexibility, we establish a framework for developing precision immunotherapies that go beyond vaccination to treat autoimmune and inflammatory diseases. These results not only expand the functional scope of mRNA-LNPs but also contribute to a growing body of work advancing antigen-specific tolerizing strategies as a new frontier in immune modulation.

## MATERIALS AND METHODS

### In Vitro Transcribed mRNA Production

Plasmids encoding codon-optimized genes of interest flanked between optimized untranslated regions and containing a 101-base poly(A) tail were supplied by GenScript (Piscataway, NJ, USA) as 1 mg/ml industrial-grade, endotoxin-free plasmid in nuclease-free water. Plasmids were linearized, purified and subject to *in vitro* transcription using MEGAscript T7 transcription kits (Thermo Fisher Scientific, Waltham, MA, USA) using co-transcriptional capping with CleanCap (TriLink Biotechnologies, San Diego, CA, USA). Uridines were fully substituted with N1-methylpseudouridine (TriLink Biotechnologies).^23–25,78^ mRNAs were purified by cellulose chromatography to remove double stranded contaminants.^79^

### Lipid Nanoparticle Formulation

The ionizable lipids SM-102, ALC-0315, 306O_i10_, MC3, and 4A3-SC8 were obtained from BroadPharm (San Diego, CA, USA) or Echelon Biosciences (Salt Lake City, UT, USA). DOPS, BMP, DOPG, DSPC, and DMG-PEG-2000 were procured from Avanti Polar Lipids (Alabaster, AL, USA), while cholesterol and sodium citrate was sourced from Sigma Aldrich (St. Louis, MO, USA). LNPs were formulated by rapid hand mixing or by microfluidic mixing using the Nanoassemblr Ignite system (Cytiva, Marlborough, MA, USA).^80^ Briefly, lipids were diluted in ethanol and combined with mRNA diluted in citrate buffer, pH 3-4 in a 1:2 volumetric ratio for hand mixing or in a 1:3 volumetric ratio for microfluidic mixing. LNPs were dialyzed against PBS to remove residual ethanol, then characterized using dynamic light scattering (DLS, ZetaSizer, Malvern Pananalytical, Westborough, MA, USA) for size and zeta potential and using a RiboGreen assay (Thermo Fisher Scientific) for mRNA encapsulation.

### Animal Experiments

All animal protocols were approved by the Institutional Animal Care and Use Committee of the University of Pennsylvania (803941), and animal procedures were performed in accordance with the Guidelines for Care and Use of Laboratory Animals. Female C57BL/6 mice, 6-10 weeks of age, were procured from Jackson Laboratory (Bar Harbor, ME, USA) and housed in a specific pathogen-free animal facility maintained at 22 ± 2°C and 40-70% humidity with 12 h light/dark cycles. mRNA-LNPs were administered to mice intravenously via the retro-orbital plexus under anesthesia.

#### Luciferase Imaging

To assess luciferase expression in major organs, mice were intravenously injected with 5 μg of luciferase mRNA–LNP. Six hours post-injection, mice received intraperitoneal (IP) injections of 130 μL D-luciferin (30 mg/mL; Regis Technologies, Morton Grove, IL, USA). Ten minutes later, mice were sacrificed, and major organs were harvested and imaged for luminescence using an IVIS Lumina III system (Revvity, Waltham, MA, USA).

#### GFP Expression and Cytokine Analysis

To evaluate GFP expression and markers of adjuvant activity, mice were administered 10 μg of GFP mRNA–LNP via intravenous injection. Six hours post-injection, blood was collected via the submandibular vein for serum cytokine analysis. At 18 hours post-injection, mice were sacrificed, spleens were harvested, and splenocytes were processed into single-cell suspensions for flow cytometry analysis, as described below. Serum cytokines were evaluated using a Luminex assay as described below.

### Cytokine and Chemokine Quantification using Luminex

Mouse cytokine and chemokine quantification was performed using a pre-mixed commercial kit (Th1/Th2/Th9/Th17/Th22/Treg Cytokine 17-Plex Mouse ProcartaPlex™ Panel: EPX170-26087-901, Thermo Fisher Scientific). Sera were collected from mice 6 hours after LNP treatment as described above, aliquoted and frozen at −80C, and thawed at the time of cytokine analysis. Assays were established per manufacturer recommendations. Data were acquired on a FlexMAP 3D quantification instrument, and analysis was done using Luminex® xPONENT® 4.2 and Bio-Plex Manager™ Software 6.1. Data quality was determined by ensuring the standard curve for each analyte had a 5P R2 value > 0.95 with or without minor fitting using xPONENT software.

### Experimental Autoimmune Encephalomyelitis

EAE induction was performed using the Hooke Laboratories MOG_35-55_ EAE induction kit (EK-2110, Lawrence, MA, USA). 10-week old, female C57BL/6 mice were injected with 100 μL MOG_35-55_/CFA emulsion SQ in two sites over the back. Mice were subsequently IP injected with 100 ng PTX 2 and 24 h after the emulsion. EAE-induced mice were immunized intravenously via retro-orbital injection with 1 μg OVA or MOG_35-55_-encoding LNPs (or equivalent volume saline) on days 7 and 10 post-EAE induction. Mice were scored by blinded observers beginning on Day 11 according to the rubric set by Hooke Laboratories (**Fig S4**). At a score of 2.5 or higher, mice were provided with wet food, and animals that scored 4 for two days in a row were considered to have reached their humane endpoint, euthanized, and assigned a score of 5 for the remainder of the study.

### Flow Cytometric Analysis

#### Splenocyte Isolation and Flow Cytometry

Spleens were processed into single-cell suspensions by mechanical dissociation using a syringe plunger through a 70 μm cell strainer. Cells were pelleted by centrifugation, resuspended in ACK Lysing Buffer (Thermo Fisher Scientific), and incubated at room temperature for 5 minutes to lyse red blood cells. Following lysis, cells were washed with FACS buffer (PBS containing 0.5% BSA and 2 mM EDTA), counted, and 5 × 10⁶ cells per sample were used for staining. Viability was assessed using LIVE/DEAD Fixable Near-IR stain (Thermo Fisher Scientific) diluted 1:2000 in PBS and incubated for 30 minutes at 37 °C, protected from light. After washing with FACS buffer, cells were stained with extracellular antibodies (**Fig S6**) diluted in FACS buffer containing 1:500 anti-mouse TruStain FcX (BioLegend, San Diego, CA, USA) for 30 minutes at 4 °C, protected from light. Stained cells were washed twice and analyzed immediately using an LSRFortessa flow cytometer (BD Biosciences, Franklin Lakes, NJ, USA).

#### Spinal Cord Dissociation and Flow Cytometry

At peak disease (∼day 18 post-induction), mice were transcardially perfused with 1x PBS to remove blood from tissue. Spinal cords were extruded as previously described.^81^ Spinal cords were dissociated according to the protocol “Dissociation of inflamed neural tissue using the Multi Dissociation Kit 1” (Miltenyi Biotec, Inc, Auburn, CA, USA). Briefly, two to three spinal cords were pooled and mechanically dissociated before being incubated with the enzyme mix for further dissociation using a Gentle MACS^TM^ Dissociator (Miltenyi Biotec, Inc, Auburn, CA, USA). Dissociated spinal cords were then filtered (70 μm), washed, and layered onto a debris removal buffer to remove myelin and other debris. The resulting single cell pellet was washed with FACS buffer (PBS containing 0.5% BSA and 2 mM EDTA) and first stained with tetramers obtained from the NIH Tetramer Core Facility that bind mouse MOG_38-49_ presented on I-Ab. Tetramer staining was performed with both PE and APC-labeled tetramers diluted 1:100 in FACS buffer containing 1:500 anti-mouse FcX for one hour at 37°C, protected from light. Cells that appeared double positive for both PE and APC were considered tetramer+ by flow cytometry. Tetramer-stained cells were subsequently stained with LIVE/DEAD Fixable NIR as described above, then fixed and permeabilized using a Foxp3 Staining Buffer Set (Thermo Fisher) for 1 hour at 4°C, protected from light. Cells were subsequently stained with antibodies (**Fig S6**) diluted in FACS buffer containing 1:500 anti-mouse TruStain FcX for 1 hour at 4°C. Samples were analyzed using a BD FACSymphony A5 SE. Flow cytometry data were collected using FACSDiva and analyzed using FlowJo software.

#### Immunohistochemistry

At peak disease, animals were perfused with ice cold 1x PBS followed by 10% formalin. Spinal cords were dissected, post-fixed overnight in 10% formalin, and left in 70% ethanol before embedding the tissue in paraffin. 10 μm sections were stained for myelin using MBP as a marker. After 3 xylene incubations of 5 minutes, tissues were rehydrated in an ethanol series from 100% to 50%, with 3 minutes incubation in each solution. Tissues were transferred to water before a 10-minute incubation in 3% H_2_O_2_ to block endogenous peroxidase activity. After a 30 minutes incubation in blocking buffer (1x PBS with 3% BSA and 5% goat serum), the primary antibody (MBP, anti-mouse, MAB382, Millipore, Burlington, MA) was diluted in 1x PBS and incubated overnight at 4°C. The next day, 3 washes in 1x PBS were performed before adding the secondary antibody (Goat anti-mouse – HRP, MP-7452, VectorLabs, Burlingame, CA) for 1h at room temperature. Another 3 washes in 1x PBS were performed before DAB development (SK-4100, vectorLabs, Burlingame, CA) for 30 seconds to 1 minute. A water wash was followed by counterstaining in hematoxylin QS (H-3404-100, VectorLabs, Burlingame, CA) for 20-30 seconds and a running water wash. Tissues were dehydrated in the ethanol series with a final 3 minute xylene incubation before mounting the tissue in VectaMount Express Mounting Media (H-5700-60, VectorLabs, Burlingame, CA). All other tissues (lungs, heart, kidney, liver and spleen) were processed by the Comparative Pathology Core at the University of Pennsylvania for embedding in paraffin, sectioning and H&E staining. All tissues were imaged on an Axio Observer 7 (Zeiss, Oberhochen, DE).

#### Statistical Analysis and Schematics

All data were imported into GraphPad Prism (Dotmatics, Boston, MA, UA) for statistical analysis. Comparisons between three or more groups were evaluated by one-way ANOVA, with p<0.05 taken to indicate statistical significance. Linear correlations were assessed using Pearson’s correlation coefficient, where values less than 0.3 were considered weak correlations, values between 0.3-0.49 were considered moderate, and values 0.5 or greater were considered strong. Schematics and cartoons were generated using BioRender.

## ACKNOWLEDGMENTS

We acknowledge and thank the Penn Cytomics and Cell Sorting Core (RRID: SCR-022376), the Penn Vet Comparative Pathology Core (RRID: SCR_022438), part of the Abramson Cancer Center Support Grant (P30 CA016520), and the Human Immunology Core at the Perelman School of Medicine at the University of Pennsylvania (RRID: SCR_022380) for assistance with Luminex assays. The HIC is supported in part by NIH P30 AI045008 and P30 CA016520. J.R.M. acknowledges funding from a UPenn Institute for RNA Innovation Pilot Award and the UPenn Institute for Immunology and Immune Health Roberts Family-Katalin Karikó Fellowship in Vaccine Research. J.R.M. and D.W. acknowledge funding from Breakthrough T1D (3-SRA-2024-1612-S-B) and from the Department of Defense (HT94252510251). L.S.C. and D.W. acknowledge support from the Tang Prize (40143777). H.S. acknowledges funding from the NIH (R01-AI136945-04). R.P.T. acknowledges support from the Cell & Molecular Biology Graduate Group in the Perelman School of Medicine at the University of Pennsylvania.

## AUTHOR CONTRIBUTIONS

J.R.M.: Conceptualization, methodology, data curation, formal analysis, investigation, writing - original draft, writing - review & editing, visualization, supervision, funding acquisition, project administration, validation; J.M.R.: Data curation, investigation, writing - review & editing; M.K.: Data curation, investigation, writing - review & editing; L.S.C.: Methodology, data curation, investigation, writing - review & editing, visualization, supervision, funding acquisition, project administration, validation; R.P.T.: Data curation, investigation, writing - review & editing; N.S.B.: Data curation, investigation, writing - review & editing; H.N.: Data curation, investigation, writing - review & editing; H.S.: Methodology, data curation, investigation, writing - review & editing; D.W.: Conceptualization, writing - review & editing, supervision, funding acquisition, resources.

## REFERENCES

1. Miao, L. et al. Delivery of mRNA vaccines with heterocyclic lipids increases anti-tumor efficacy by STING-mediated immune cell activation. Nat Biotechnol 37, 1174–1185 (2019).

2. Han, X. et al. Adjuvant lipidoid-substituted lipid nanoparticles augment the immunogenicity of SARS-CoV-2 mRNA vaccines. Nat. Nanotechnol. 18, 1105–1114 (2023).

3. Hajj, K. A. & Whitehead, K. A. Tools for translation: non-viral materials for therapeutic mRNA delivery. Nat Rev Mater 2, 17056 (2017).

4. Pardi, N. et al. Nucleoside-modified mRNA immunization elicits influenza virus hemagglutinin stalk-specific antibodies. Nat Commun 9, 3361 (2018).

5. Ackun-Farmmer, M. A. & Jewell, C. M. Delivery Route Considerations for Designing Antigen-Specific Biomaterial Strategies to Combat Autoimmunity. Advanced NanoBiomed Research 3, 2200135 (2023).

6. Breda, L. et al. In vivo hematopoietic stem cell modification by mRNA delivery. Science 381, 436–443 (2023).

7. Ni, H. et al. Piperazine-derived lipid nanoparticles deliver mRNA to immune cells in vivo. Nat Commun 13, 4766 (2022).

8. Melamed, J. R. et al. Lipid nanoparticle chemistry determines how nucleoside base modifications alter mRNA delivery. Journal of Controlled Release 341, 206–214 (2022).

9. Chen, J. et al. Lipid nanoparticle-mediated lymph node–targeting delivery of mRNA cancer vaccine elicits robust CD8 ^+^ T cell response. Proc. Natl. Acad. Sci. U.S.A. 119, e2207841119 (2022).

10. Fenton, O. S. et al. Synthesis and Biological Evaluation of Ionizable Lipid Materials for the In Vivo Delivery of Messenger RNA to B Lymphocytes. Advanced Materials 29, 1606944 (2017).

11. Razavi, R., Kegel, M., Muscat-Rivera, J., Weissman, D. & Melamed, J. R. Harnessing mRNA-lipid nanoparticles as innovative therapies for autoimmune diseases. Molecular Therapy Methods & Clinical Development 33, 101566 (2025).

12. Tombácz, I. et al. Highly efficient CD4+ T cell targeting and genetic recombination using engineered CD4+ cell-homing mRNA-LNPs. Molecular Therapy 29, 3293–3304 (2021).

13. Rurik, J. G., et al. CAR T cells produced in vivo to treat cardiac injury. (2022).

14. Billingsley, M. M. et al. In Vivo mRNA CAR T Cell Engineering via Targeted Ionizable Lipid Nanoparticles with Extrahepatic Tropism. Small 2304378 (2023) doi:10.1002/smll.202304378.

15. Cheng, Q. et al. Selective organ targeting (SORT) nanoparticles for tissue-specific mRNA delivery and CRISPR–Cas gene editing. Nat. Nanotechnol. 15, 313–320 (2020).

16. Dilliard, S. A., Cheng, Q. & Siegwart, D. J. On the mechanism of tissue-specific mRNA delivery by selective organ targeting nanoparticles. Proc. Natl. Acad. Sci. U.S.A. 118, e2109256118 (2021).

17. LoPresti, S. T., Arral, M. L., Chaudhary, N. & Whitehead, K. A. The replacement of helper lipids with charged alternatives in lipid nanoparticles facilitates targeted mRNA delivery to the spleen and lungs. Journal of Controlled Release 345, 819–831 (2022).

18. Luozhong, S. et al. Phosphatidylserine Lipid Nanoparticles Promote Systemic RNA Delivery to Secondary Lymphoid Organs. Nano Lett. 22, 8304–8311 (2022).

19. Álvarez-Benedicto, E. et al. Spleen SORT LNP Generated in situ CAR T Cells Extend Survival in a Mouse Model of Lymphoreplete B Cell Lymphoma. Angew Chem Int Ed Engl 62, e202310395 (2023).

20. Kranz, L. M. et al. Systemic RNA delivery to dendritic cells exploits antiviral defence for cancer immunotherapy. Nature 534, 396–401 (2016).

21. Krienke, C. et al. A noninflammatory mRNA vaccine for treatment of experimental autoimmune encephalomyelitis. Science 371, 145–153 (2021).

22. Grunwitz, C. et al. HPV16 RNA-LPX vaccine mediates complete regression of aggressively growing HPV-positive mouse tumors and establishes protective T cell memory. OncoImmunology 8, e1629259 (2019).

23. Karikó, K., Buckstein, M., Ni, H. & Weissman, D. Suppression of RNA Recognition by Toll-like Receptors: The Impact of Nucleoside Modification and the Evolutionary Origin of RNA. Immunity 23, 165–175 (2005).

24. Karikó, K. et al. Incorporation of Pseudouridine Into mRNA Yields Superior Nonimmunogenic Vector With Increased Translational Capacity and Biological Stability. Molecular Therapy 16, 1833–1840 (2008).

25. Andries, O. et al. N1-methylpseudouridine-incorporated mRNA outperforms pseudouridine-incorporated mRNA by providing enhanced protein expression and reduced immunogenicity in mammalian cell lines and mice. Journal of Controlled Release 217, 337–344 (2015).

26. Sittplangkoon, C. et al. mRNA vaccine with unmodified uridine induces robust type I interferon-dependent anti-tumor immunity in a melanoma model. Front. Immunol. 13, 983000 (2022).

27. Pardi, N. et al. Nucleoside-modified mRNA vaccines induce potent T follicular helper and germinal center B cell responses. Journal of Experimental Medicine 215, 1571–1588 (2018).

28. Alameh, M.-G. et al. Lipid nanoparticles enhance the efficacy of mRNA and protein subunit vaccines by inducing robust T follicular helper cell and humoral responses. Immunity 54, 2877–2892.e7 (2021).

29. Parhiz, H. et al. Added to pre-existing inflammation, mRNA-lipid nanoparticles induce inflammation exacerbation (IE). Journal of Controlled Release 344, 50–61 (2022).

30. Baden, L. R. et al. Efficacy and Safety of the mRNA-1273 SARS-CoV-2 Vaccine. N Engl J Med 384, 403–416 (2021).

31. Polack, F. P. et al. Safety and Efficacy of the BNT162b2 mRNA Covid-19 Vaccine. N Engl J Med 383, 2603–2615 (2020).

32. Adams, D. et al. Patisiran, an RNAi Therapeutic, for Hereditary Transthyretin Amyloidosis. N Engl J Med 379, 11–21 (2018).

33. Hassett, K. J. et al. Optimization of Lipid Nanoparticles for Intramuscular Administration of mRNA Vaccines. Mol Ther Nucleic Acids 15, 1–11 (2019).

34. Hajj, K. A. et al. A Potent Branched-Tail Lipid Nanoparticle Enables Multiplexed mRNA Delivery and Gene Editing *In Vivo*. Nano Lett. 20, 5167–5175 (2020).

35. Hajj, K. A. et al. Branched-Tail Lipid Nanoparticles Potently Deliver mRNA In Vivo due to Enhanced Ionization at Endosomal pH. Small 15, e1805097 (2019).

36. Melamed, J. R. et al. Ionizable lipid nanoparticles deliver mRNA to pancreatic β cells via macrophage-mediated gene transfer. Sci. Adv. 9, eade1444 (2023).

37. Den Haan, J. M. M., Lehar, S. M. & Bevan, M. J. Cd8+ but Not Cd8− Dendritic Cells Cross-Prime Cytotoxic T Cells in Vivo. The Journal of Experimental Medicine 192, 1685–1696 (2000).

38. Hildner, K., et al. *Batf3* Deficiency Reveals a Critical Role for CD8α^+^ Dendritic Cells in Cytotoxic T Cell Immunity. Science 322, 1097–1100 (2008).

39. Lyons-Cohen, M. R., Shamskhou, E. A. & Gerner, M. Y. Site-specific regulation of Th2 differentiation within lymph node microenvironments. Journal of Experimental Medicine 221, e20231282 (2024).

40. Ngo, C., Garrec, C., Tomasello, E. & Dalod, M. The role of plasmacytoid dendritic cells (pDCs) in immunity during viral infections and beyond. Cell Mol Immunol 21, 1008–1035 (2024).

41. Lenschow, D. J., Walunas, T. L. & Bluestone, J. A. CD28/B7 SYSTEM OF T CELL COSTIMULATION. Annu. Rev. Immunol. 14, 233–258 (1996).

42. Maj, T., Slawek, A. & Chelmonska-Soyta, A. CD80 and CD86 costimulatory molecules differentially regulate OT-II CD4^+^ T lymphocyte proliferation and cytokine response in cocultures with antigen-presenting cells derived from pregnant and pseudopregnant mice. Mediators Inflamm 2014, 769239 (2014).

43. Grewal, I. S. & Flavell, R. A. The role of CD40 ligand in costimulation and T-cell activation. Immunol Rev 153, 85–106 (1996).

44. Peng, X., Kasran, A., Warmerdam, P. A., de Boer, M. & Ceuppens, J. L. Accessory signaling by CD40 for T cell activation: induction of Th1 and Th2 cytokines and synergy with interleukin-12 for interferon-gamma production. Eur J Immunol 26, 1621–1627 (1996).

45. Machen, J. et al. Antisense oligonucleotides down-regulating costimulation confer diabetes-preventive properties to nonobese diabetic mouse dendritic cells. J Immunol 173, 4331–4341 (2004).

46. Di Caro, V., Giannoukakis, N. & Trucco, M. In vivo delivery of nucleic acid-formulated microparticles as a potential tolerogenic vaccine for type 1 diabetes. Rev Diabet Stud 9, 348–356 (2012).

47. Suzuki, M., Yokota, M., Matsumoto, T. & Ozaki, S. Synergic Effects of CD40 and CD86 Silencing in Dendritic Cells on the Control of Allergic Diseases. Int Arch Allergy Immunol 177, 87–96 (2018).

48. Zhang, Q. et al. Permanent acceptance of mouse cardiac allografts with CD40 siRNA to induce regulatory myeloid cells by use of a novel polysaccharide siRNA delivery system. Gene Ther 22, 217–226 (2015).

49. Constantinescu, C. S., Farooqi, N., O’Brien, K. & Gran, B. Experimental autoimmune encephalomyelitis (EAE) as a model for multiple sclerosis (MS). Br J Pharmacol 164, 1079–1106 (2011).

50. Jakimovski, D. et al. Multiple sclerosis. The Lancet 403, 183–202 (2024).

51. Ciccarelli, O. et al. Pathogenesis of multiple sclerosis: insights from molecular and metabolic imaging. Lancet Neurol 13, 807–822 (2014).

52. Absinta, M. et al. A lymphocyte-microglia-astrocyte axis in chronic active multiple sclerosis. Nature 597, 709–714 (2021).

53. Hauser, S. L. & Cree, B. A. C. Treatment of Multiple Sclerosis: A Review. Am J Med 133, 1380–1390.e2 (2020).

54. Mendel, I., De Rosbo, N. K. & Ben-Nun, A. A myelin oligodendrocyte glycoprotein peptide induces typical chronic experimental autoimmune encephalomyelitis in H-2^b^ mice: Fine specificity and T cell receptor Vβ expression of encephalitogenic T cells. Eur J Immunol 25, 1951–1959 (1995).

55. El-behi, M., Rostami, A. & Ciric, B. Current views on the roles of Th1 and Th17 cells in experimental autoimmune encephalomyelitis. J Neuroimmune Pharmacol 5, 189–197 (2010).

56. Hunter, Z. et al. A Biodegradable Nanoparticle Platform for the Induction of Antigen-Specific Immune Tolerance for Treatment of Autoimmune Disease. ACS Nano 8, 2148–2160 (2014).

57. Chen, N. et al. Co-Delivery of Disease Associated Peptide and Rapamycin via Acetalated Dextran Microparticles for Treatment of Multiple Sclerosis. Adv. Biosys. 1, 1700022 (2017).

58. Rhodes, K. R. et al. Bioengineered particles expand myelin-specific regulatory T cells and reverse autoreactivity in a mouse model of multiple sclerosis. Sci. Adv. 9, eadd8693 (2023).

59. Yeste, A., Nadeau, M., Burns, E. J., Weiner, H. L. & Quintana, F. J. Nanoparticle-mediated codelivery of myelin antigen and a tolerogenic small molecule suppresses experimental autoimmune encephalomyelitis. Proc Natl Acad Sci U S A 109, 11270–11275 (2012).

60. Kwiatkowski, A. J. et al. Treatment with an antigen-specific dual microparticle system reverses advanced multiple sclerosis in mice. Proc. Natl. Acad. Sci. U.S.A. 119, e2205417119 (2022).

61. Peine, K. J. et al. Treatment of Experimental Autoimmune Encephalomyelitis by Codelivery of Disease Associated Peptide and Dexamethasone in Acetalated Dextran Microparticles. *Mol*. Pharmaceutics 11, 828–835 (2014).

62. Lewis, J. S. et al. Dual-Sized Microparticle System for Generating Suppressive Dendritic Cells Prevents and Reverses Type 1 Diabetes in the Nonobese Diabetic Mouse Model. ACS Biomater Sci Eng 5, 2631–2646 (2019).

63. Lewis, J. S. et al. A combination dual-sized microparticle system modulates dendritic cells and prevents type 1 diabetes in prediabetic NOD mice. Clinical Immunology 160, 90–102 (2015).

64. Chen, N., Kroger, C. J., Tisch, R. M., Bachelder, E. M. & Ainslie, K. M. Prevention of Type 1 Diabetes with Acetalated Dextran Microparticles Containing Rapamycin and Pancreatic Peptide P31. Adv Healthcare Materials 7, 1800341 (2018).

65. Daniel, C., Weigmann, B., Bronson, R. & Von Boehmer, H. Prevention of type 1 diabetes in mice by tolerogenic vaccination with a strong agonist insulin mimetope. Journal of Experimental Medicine 208, 1501–1510 (2011).

66. Prasad, S. et al. Tolerogenic Ag-PLG nanoparticles induce tregs to suppress activated diabetogenic CD4 and CD8 T cells. J Autoimmun 89, 112–124 (2018).

67. Freitag, T. L. et al. Gliadin Nanoparticles Induce Immune Tolerance to Gliadin in Mouse Models of Celiac Disease. Gastroenterology 158, 1667–1681.e12 (2020).

68. Benne, N. et al. Autoantigen-Dexamethasone Conjugate-Loaded Liposomes Halt Arthritis Development in Mice. Adv Healthc Mater 13, e2304238 (2024).

69. Ter Braake, D., Benne, N., Lau, C. Y. J., Mastrobattista, E. & Broere, F. Retinoic Acid-Containing Liposomes for the Induction of Antigen-Specific Regulatory T Cells as a Treatment for Autoimmune Diseases. Pharmaceutics 13, 1949 (2021).

70. Thatte, A. S., Billingsley, M. M., Weissman, D., Melamed, J. R. & Mitchell, M. J. Emerging strategies for nanomedicine in autoimmunity. Adv Drug Deliv Rev 207, 115194 (2024).

71. Gomi, M. et al. Tolerogenic Lipid Nanoparticles for Delivering Self-Antigen mRNA for the Treatment of Experimental Autoimmune Encephalomyelitis. Pharmaceuticals (Basel) 16, 1270 (2023).

72. Benne, N., Ter Braake, D., Stoppelenburg, A. J. & Broere, F. Nanoparticles for Inducing Antigen-Specific T Cell Tolerance in Autoimmune Diseases. Front Immunol 13, 864403 (2022).

73. Ramos, G. C. et al. Apoptotic mimicry: phosphatidylserine liposomes reduce inflammation through activation of peroxisome proliferator-activated receptors (PPARs) in vivo. Br J Pharmacol 151, 844–850 (2007).

74. Nagata, S., Hanayama, R. & Kawane, K. Autoimmunity and the clearance of dead cells. Cell 140, 619–630 (2010).

75. Villalba, A. et al. Preclinical evaluation of antigen-specific nanotherapy based on phosphatidylserine-liposomes for type 1 diabetes. Artif Cells Nanomed Biotechnol 48, 77–83 (2020).

76. Rodriguez-Fernandez, S. et al. Phosphatidylserine-Liposomes Promote Tolerogenic Features on Dendritic Cells in Human Type 1 Diabetes by Apoptotic Mimicry. Front Immunol 9, 253 (2018).

77. Benne, N. et al. Anionic 1,2-distearoyl-sn-glycero-3-phosphoglycerol (DSPG) liposomes induce antigen-specific regulatory T cells and prevent atherosclerosis in mice. J Control Release 291, 135–146 (2018).

78. Karikó, K., Muramatsu, H., Keller, J. M. & Weissman, D. Increased erythropoiesis in mice injected with submicrogram quantities of pseudouridine-containing mRNA encoding erythropoietin. Mol Ther 20, 948–953 (2012).

79. Baiersdörfer, M. et al. A Facile Method for the Removal of dsRNA Contaminant from In Vitro-Transcribed mRNA. Molecular Therapy - Nucleic Acids 15, 26–35 (2019).

80. Wang, X. et al. Preparation of selective organ-targeting (SORT) lipid nanoparticles (LNPs) using multiple technical methods for tissue-specific mRNA delivery. Nat Protoc 18, 265–291 (2023).

81. Richner, M., Jager, S. B., Siupka, P. & Vaegter, C. B. Hydraulic Extrusion of the Spinal Cord and Isolation of Dorsal Root Ganglia in Rodents. J Vis Exp 55226 (2017) doi:10.3791/55226.

