## Supplemental Information for "Anionic lipids modulate mRNA-lipid nanoparticle immunogenicity and confer protection in a mouse model of multiple sclerosis"

**AFFILIATIONS**

**CORRESPONDING AUTHORS**

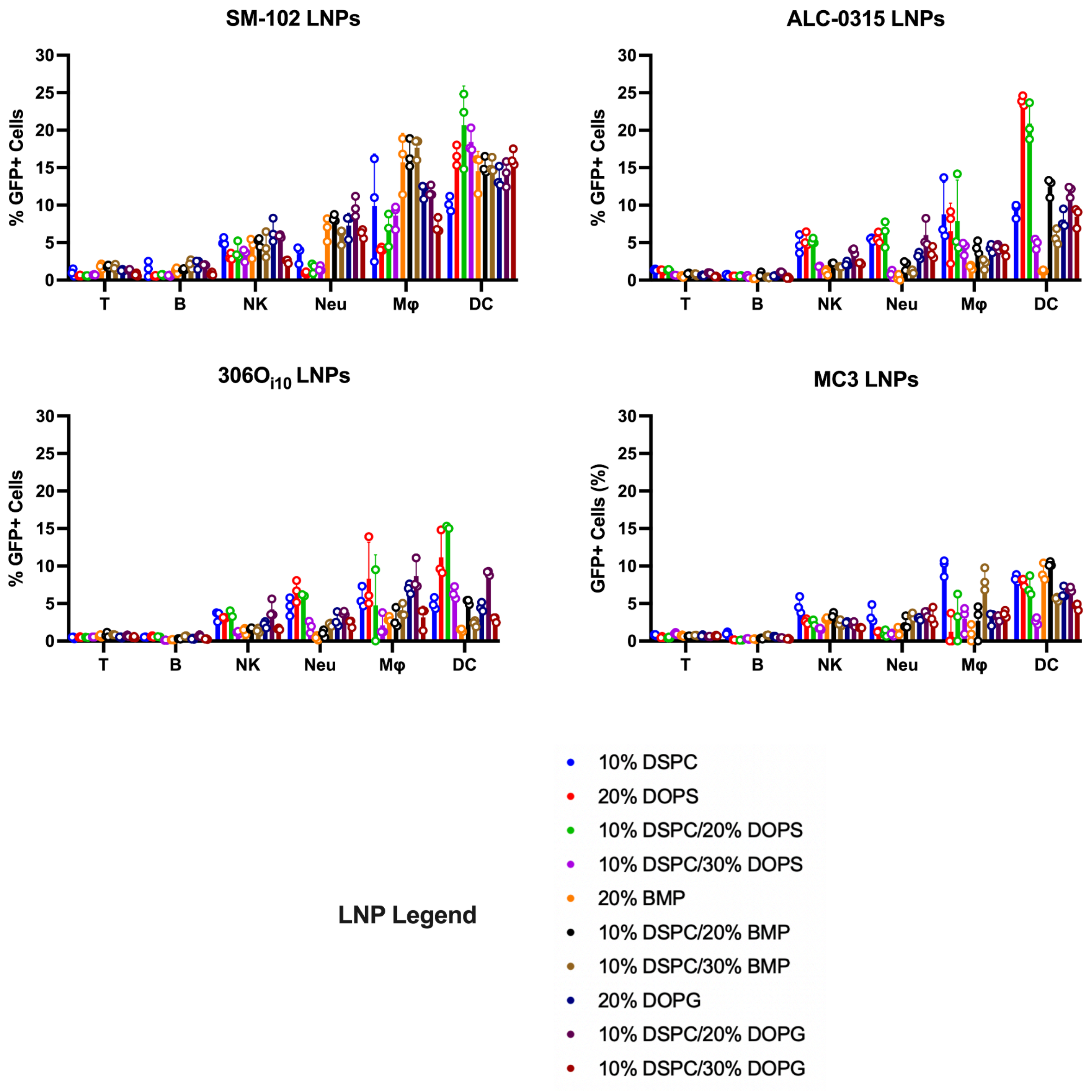


**Figure S1: Summarized flow cytometry data showing GFP expression in all splenocyte subtypes evaluated resulting from all LNP formulations tested.** C57BL/6 mice were IV injected with 10 µg GFP mRNA-LNPs. Splenocytes were harvested and evaluated by flow cytometry 24 hours later. Data shown represent n=3 mice per LNP formulation. Error bars represent standard deviations.

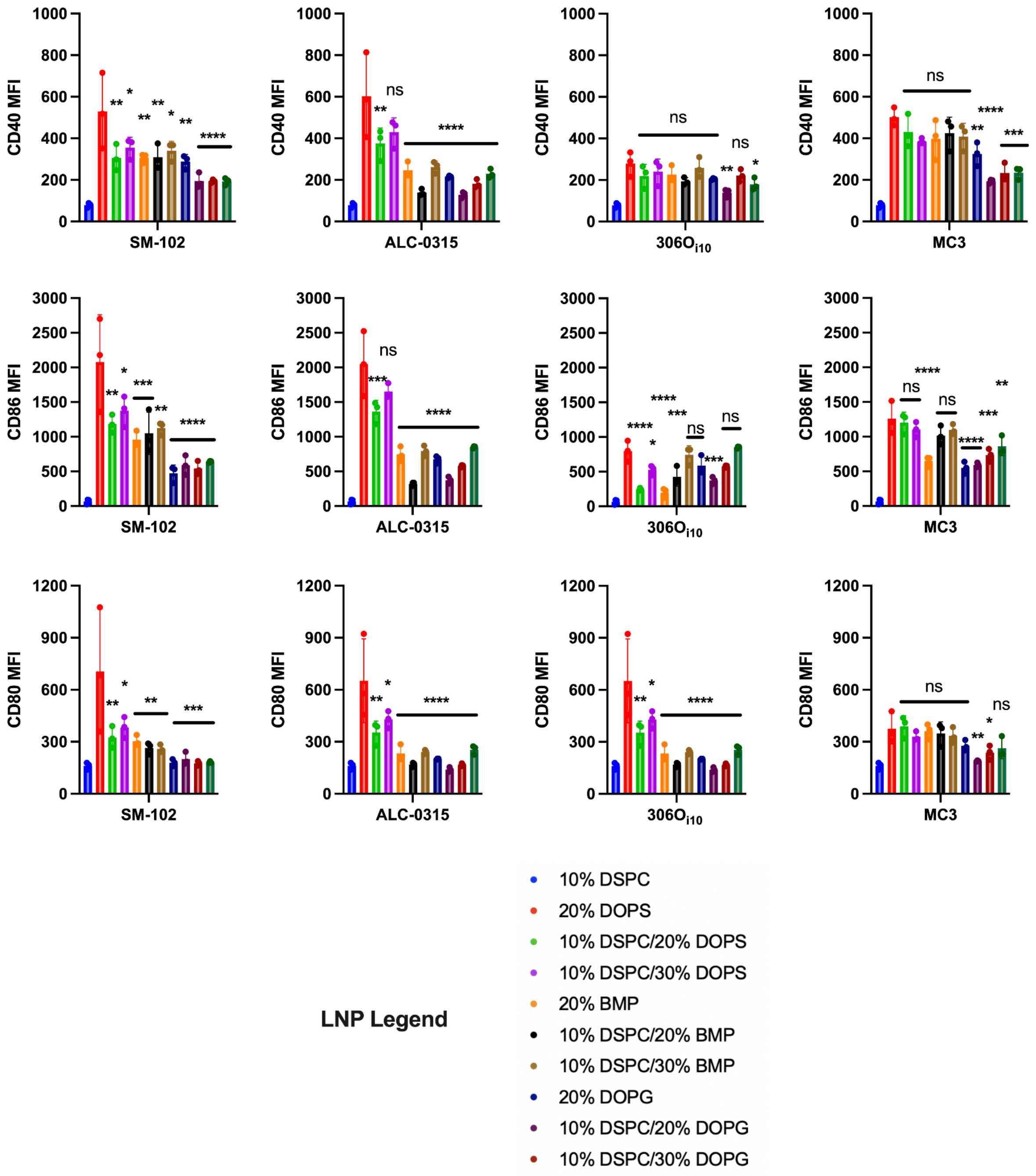


**Figure S2: Summarized flow cytometry data showing expression levels of DC activation resulting from all LNP formulations tested.** C57BL/6 mice were IV injected with 10 µg GFP mRNA-LNPs. Splenocytes were harvested and evaluated by flow cytometry 24 hours later. Data shown represent n=3 mice per LNP formulation. Error bars represent standard deviations. Statistical differences were calculated by ANOVA with post-hoc Tukey test. *$p<0.05$, **$p<0.01$, ***$p<0.005$, ****$p<0.001$ relative to 10% DSPC controls.

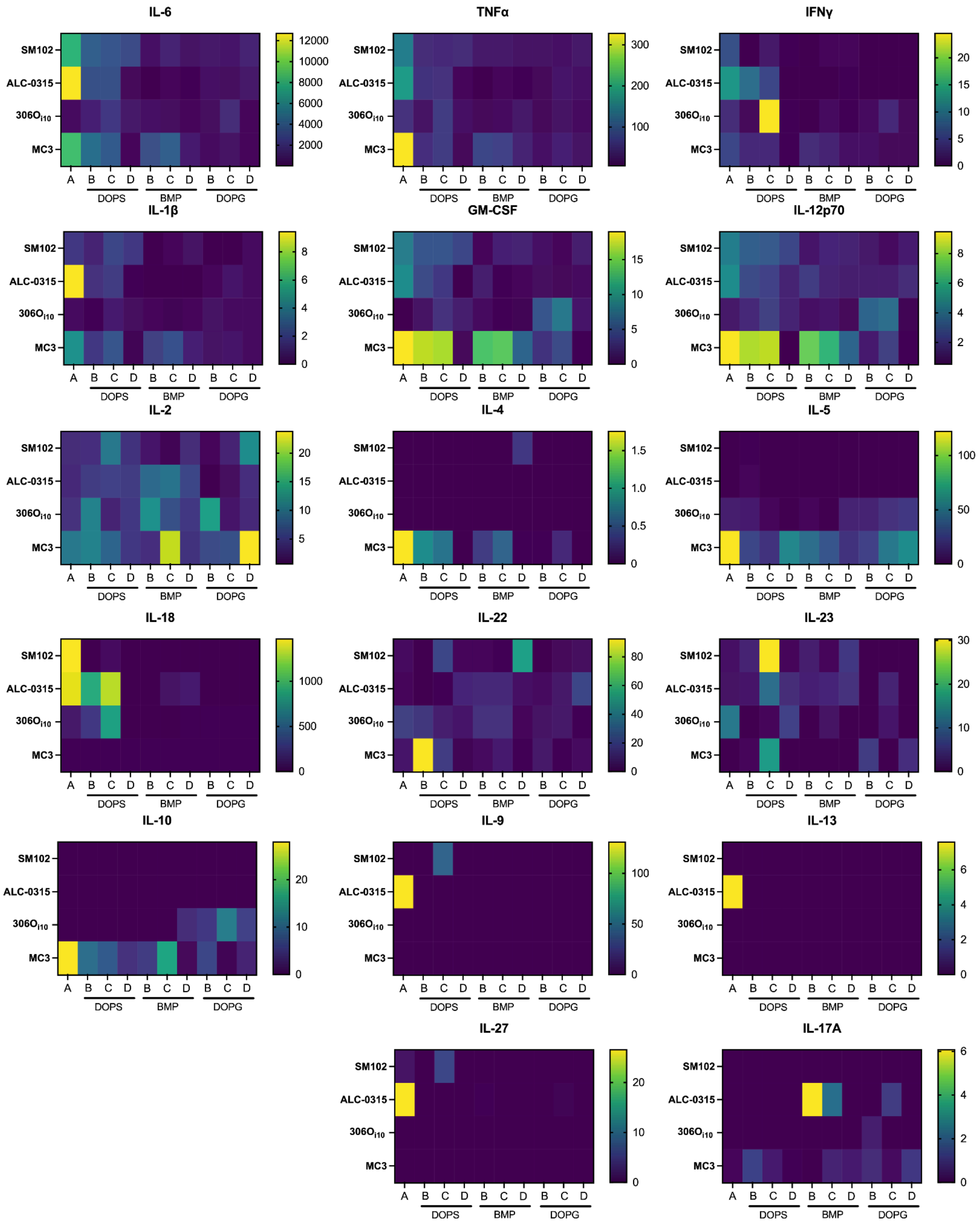


**Figure S3: All Luminex assay results for all LNPs tested and cytokines evaluated.** Heat map legends are serum cytokine concentrations in pg/mL.

| Score | Clinical observations |
|---|---|
| 0.0 | No obvious changes in motor function compared to non-immunized mice.<br><br>When picked up by base of tail, the tail has tension and is erect. Hind legs are usually spread apart. When the mouse is walking, there is no gait or head tilting. |
| 0.5 | Tip of tail is limp.<br><br>When picked up by base of tail, the tail has tension except for the tip. Muscle straining is felt in the tail, while the tail continues to move. |
| 1.0 | Limp tail.<br><br>When picked up by base of tail, instead of being erect, the whole tail drapes over finger. Hind legs are usually spread apart. No signs of tail movement are observed. |
| 1.5 | Limp tail and hind leg inhibition.<br><br>When picked up by base of tail, the whole tail drapes over finger. When the mouse is dropped on a wire rack, at least one hind leg falls through consistently. Walking is very slightly wobbly. |
| 2.0 | Limp tail and weakness of hind legs.<br><br>When picked up by base of tail, the legs are not spread apart, but held closer together. When the mouse is observed walking, it has a clearly apparent wobbly walk. One foot may have toes dragging, but the other leg has no apparent inhibitions of movement.<br><br>- OR -<br><br>Mouse appears to be at score 0.0, but there are obvious signs of head tilting when the walk is observed. The balance is poor. |
| 2.5 | Limp tail and dragging of hind legs.<br><br>Both hind legs have some movement, but both are dragging at the feet (mouse trips on hind feet).<br><br>- OR -<br><br>No movement in one leg/completely dragging one leg, but movement in the other leg.<br><br>- OR -<br><br>EAE severity appears mild when picked up (as score 0.0-1.5), but there is a strong head tilt that causes the mouse to occasionally fall over. |
| 3.0 | Limp tail and complete paralysis of hind legs (most common).<br><br>- OR -<br><br>Limp tail and almost complete paralysis of hind legs. One or both hind legs are able to paddle, but neither hind leg is able to move forward of the hind hip.<br><br>- OR -<br><br>Limp tail with paralysis of one front and one hind leg. |

| | |
|---|---|
| | - OR -<br><br>ALL of:<br><br>■ Severe head tilting,<br>■ Walking only along the edges of the cage,<br>■ Pushing against the cage wall,<br>■ Spinning when picked up by base of tail. |
| 3.5 | Limp tail and complete paralysis of hind legs. In addition to:<br><br>Mouse is moving around the cage, but when placed on its side, is unable to right itself. Hind legs are together on one side of body.<br><br>- OR -<br><br>Mouse is moving around the cage, but the hind quarters are flat like a pancake, giving the appearance of a hump in the front quarters of the mouse. |
| 4.0 | Limp tail, complete hind leg and partial front leg paralysis.<br><br>Mouse is minimally moving around the cage but appears alert and feeding.<br><br>Often euthanasia is recommended after the mouse scores 4.0 for 2 days. However, with daily s.c. fluids most C57BL/6 mice may recover to 3.5 or 3.0. When the mouse is euthanized because of severe paralysis, a score of 5.0 is entered for that mouse for the rest of the experiment. |
| 4.5 | Complete hind and partial front leg paralysis, no movement around the cage. Mouse is not alert.<br><br>Mouse has minimal movement in the front legs. The mouse barely responds to contact.<br><br>Euthanasia is recommended. When the mouse is euthanized because of severe paralysis, a score of 5.0 is entered for that mouse for the rest of the experiment. |
| 5.0 | Mouse is spontaneously rolling in the cage (euthanasia is recommended).<br><br>- OR -<br><br>Mouse is found dead due to paralysis.<br><br>- OR -<br><br>Mouse is euthanized due to severe paralysis. |

**Figure S4: EAE scoring chart from Hooke Labs.** EAE scoring was performed precisely according to this rubric, provided by Hooke Labs at https://hookelabs.com/protocols/eaeAI_C57BL6.html#:~:text=Eighty%20(80)%20to%20100%25,following%206%20to%208%20days.

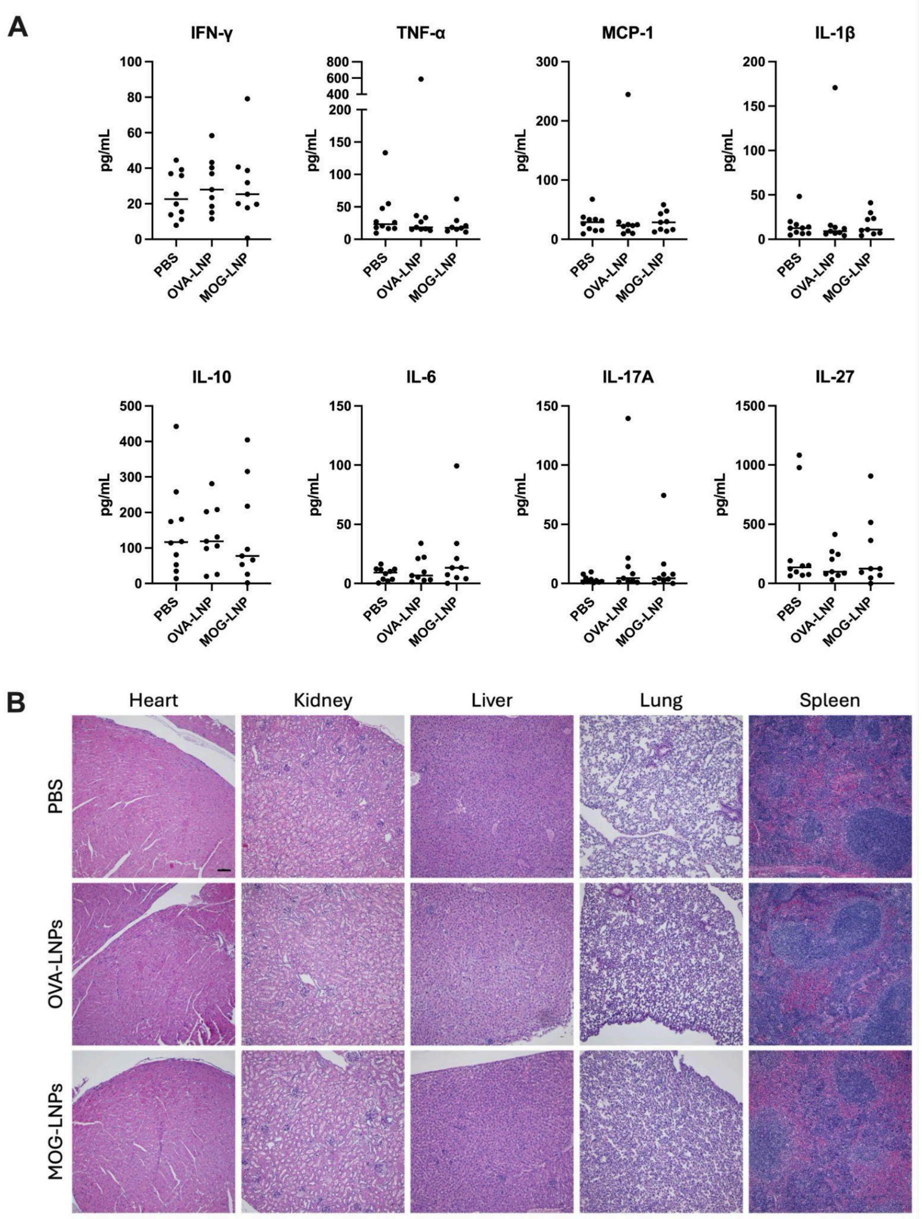


**Fig S5: LNP treatment during EAE did not induce long-term toxicity.** ***A)*** Serum was collected from mice at peak disease and analyzed for cytokine concentrations using a LegendPLEX assay. No statistical differences were observed across treatment groups, by one-way ANOVA. ***B)*** H&E staining of peripheral organs display no morphological toxicity due to LNP treatment. Heart, liver, lung, kidney and spleen were harvested from EAE mice receiving the indicated treatments at peak disease, day 18 post-EAE induction. After 1x PBS perfusion, tissues were post-fixed in 10% formalin and transferred to 70% ethanol before paraffin embedding and sectioning. Scale bar = 100 μm.

| Target | Fluorophore | Clone | Dilution | Vendor | Catalog # |
|---|---|---|---|---|---|
| **Flow cytometry panel for spleen and spinal cord (Fig.5)** | | | | | |
| CD4 | BUV395 | RM4.5 | 1:100 | BD | 568375 |
| CD8 | BUV747 | 53-6.7 | 1:100 | BD | 612759 |
| T-bet | BV421 | eBio4B10 | 1:50 | BioLegend | 644815 |
| CD25 | BV711 | PC61 | 1:50 | BioLegend | 102049 |
| CD3 | FITC | 145-2C11 | 1:100 | BioLegend | 100306 |
| RORγt | RB705 | Q31-378 | 1:50 | BD | 570259 |
| CXCR5 | RB744 | 2G8 | 1:50 | BD (OptiBuild) | 757847 |
| PD-1 | PE-Dazzle594 | RMP1-30 | 1:50 | BioLegend | 109116 |
| FoxP3 | PE-Cy7 | fjk-16s | 1:50 | ThermoFisher | 25-5775-82 |
| CD45 | BV605 | 30-F11 | 1:100 | BioLegend | 103139 |
| Viability | Live/Dead Fixable NIR | | 1:2000 | ThermoFisher | L10119 |
| **Flow cytometry panel for total splenocytes (Fig.2)** | | | | | |
| CD3 | PerCP-Cy5.5 | 17A2 | 1:100 | BioLegend | 100218 |
| CD4 | BUV395 | RM4.5 | 1:100 | BD | 568375 |
| CD8 | BUV805 | 53-6.7 | 1:100 | BD | 621898 |
| CD11b | APC-Fire750 | M1/70 | 1:100 | BioLegend | 101262 |
| CD11c | BV421 | N418 | 1:100 | BioLegend | 117330 |
| CD19 | BV605 | 6D5 | 1:100 | BioLegend | 115539 |
| CD69 | APC | H1.2F3 | 1:100 | BioLegend | 104514 |
| F4/80 | PE/Cy7 | QA17A29 | 1:50 | BioLegend | 157307 |
| Ly6C | PE | HK1.4 | 1:100 | BioLegend | 128008 |
| Ly6G | AF700 | 1A8 | 1:100 | BioLegend | 127622 |
| I-A/I-E | BV650 | M5/114.15.2 | 1:100 | BioLegend | 107641 |
| NK1.1 | BV711 | PK136 | 1:100 | BioLegend | 108745 |

| | | | | | |
|---|---|---|---|---|---|
| **Flow cytometry panel for DCS (Fig.2)** | | | | | |
| B220 | BV711 | RA3-6B2 | 1:100 | BioLegend | 103225 |
| CD8a | PerCP-Cy5.5 | 53-6.7 | 1:100 | BioLegend | 100734 |
| CD11b | APC-Fire750 | M1/70 | 1:100 | BioLegend | 101262 |
| CD11c | BV421 | N418 | 1:100 | BioLegend | 117330 |
| CD40 | PE | FGK45 | 1:100 | BioLegend | 157506 |
| CD80 | PE-Cy7 | 16-10A1 | 1:100 | BioLegend | 104734 |
| CD83 | APC | Michel-19 | 1:100 | BioLegend | 121510 |
| CD86 | BV605 | GL-1 | 1:100 | BioLegend | 105037 |
| CD19 (dump) | AF594 | 6D5 | 1:100 | BioLegend | 115552 |
| F4/80 (dump) | AF594 | BM8 | 1:50 | BioLegend | 123140 |
| Ly6G (dump) | AF594 | 1A8 | 1:100 | BioLegend | 127636 |
| I-A/I-E | BV650 | M5/114.15.2 | 1:100 | BioLegend | 107641 |
| PDCA1 | SUV387 | 927 | 1:100 | BioLegend | 127044 |
| Viability | Live/Dead Fixable Aqua | n/a | 1:2000 | ThermoFisher | L34957 |
| **Antibody for Immunohistochemistry (Fig.5D)** | | | | | |
| MBP | n/a | 129-138, clone | 1:100 | Millipore | MAB382 |

**Table S1. Antibody information.**
